# Crystal structures of Arabidopsis and Physcomitrella CR4 reveal the molecular architecture of CRINKLY4 receptor kinases

**DOI:** 10.1101/2020.08.10.245050

**Authors:** Satohiro Okuda, Ludwig A. Hothorn, Michael Hothorn

**Author notes:** Department of Biological Science, School of Science, University of Tokyo, 113-0033 Tokyo, Japan (Satohiro Okuda). retired (Ludwig A. Hothorn).

## Abstract

Plant-unique receptor kinases harbor conserved cytoplasmic kinase domains and sequence-diverse ectodomains. Here we report crystal structures of CRINKLY4-type ectodomains from Arabidopsis ACR4 and *Physcomitrella patens* PpCR4 at 1.95 Å and 2.70 Å resolution, respectively. Monomeric CRINKLY4 ectodomains harbor a N-terminal WD40 domain and a cysteine-rich domain (CRD) connected by a short linker. The WD40 domain forms a seven-bladed β-propeller with the N-terminal strand buried in its center. Each propeller blade is stabilized by a disulfide bond and contributes to the formation of a putative ligand binding groove. The CRD forms a β-sandwich structure stabilized by six disulfide bonds and shares low structural homology with tumor necrosis factor receptor domains. Quantitative binding assays reveal that ACR4 is not a direct receptor for the peptide hormone CLE40. An ACR4 variant lacking the entire CRD can rescue the known *acr4-2* mutant phenotype, as can expression of PpCR4. Together, an evolutionary conserved signaling function for CRINKLY4 receptor kinases is encoded in its WD40 domain.

## Introduction

Plants have evolved a unique set of membrane receptor kinases (RKs) that regulate diverse aspects of growth and development, form the first layer of the plant immune system and mediate symbiotic interactions. RKs contain a single membrane-spanning helix, a conserved dual-specificity cytoplasmic kinase domain and sequence-diverse extracellular domains (ectodomains) involved in signal perception and receptor activation^1^. The three-dimensional structures and functions of plant RKs with leucine-rich repeat (LRR) ectodomains have been characterized in detail, yielding a molecular understanding of their ligand binding and receptor activation mechanisms^2^.

Crystal structures of non-LRR RKs have been reported for lysine-motif domain containing immune and symbiosis receptors involved in the perception of N-acetyl-D-glucosamin-containing ligands^3–5^. S-locus receptor kinases involved in self recognition during flower pollination have been structurally characterized to contain β-barrel lectin domains and growth factor-like domains, all contributing to the specific recognition of a cysteine-rich signaling peptide^6^. Two other classes of RKs with lectin domain-containing extracellular domains have subsequently been characterized^7^: The CYSTEINE-RICH RECEPTOR-LIKE PROTEIN KINASES (CRKs) contain a tandem arrangement of DOMAIN OF UNKNOWN FUNCTION 26 (DUF26) lectin domains, which may be involved in the recognition of a carbohydrate ligand^8^. The *Catharanthus roseus* receptor kinase 1-like (CrRLK1L) family contains a tandem arrangement of malectin domains^9^ involved in the sensing of cysteine-rich RAPID ALKALINIZATION FACTOR peptides^10^, which can be distinctly bound to either LORELEI-like GLYCOLPHOSPHATIDYLINOSITOL (GPI)-ANCHORED PROTEINS^10^ or to the LRR domains of extensins^11^.

Plant-unique CRINKLY4 (CR4) -type RKs show an unusual ectodomain structure radically different from the known LRR, LysM and lectin receptor kinases described above. The founding member of this family was identified by mapping the *crinkly4* mutation affecting leaf epidermis differentiation in maize^12^. The putative receptor kinase CR4 was initially shown to contain an active cytoplasmic protein kinase module as well as an ectodomain with distant sequence homology to tumor necrosis factor receptor (TNFR) domains^12,13^. TNFR type I and II receptors contain a cysteine-rich ectodomain that folds into several ∼40 amino-acid segments. Each segment contains 6 conserved cysteines engaged in disulfide bonds^14^ and can act as binding sites for growth factors^15^. The sequence similarities between the CRINKLY4 and TNFR ectodomains suggested a role for maize CR4 in growth factor-triggered cell differentiation responses^13,16^. Anti-sense knock-down or insertion mutation-based knock-out of *ACR4*, the Arabidopsis ortholog of maize CR4, again resulted in epidermis differentiation defects, leading to, for example, abnormal embryo and seed development^17–19^. ACR4 localizes to the plasma membrane and to endosomes^17–20^ and is a catalytically active protein kinase^18,21–23^. Sequence analysis of four ACR4 homologs in Arabidopsis indicated the presence of a conserved N-terminal β-propeller structure in CRINKLY4 ectodomains^21^. Subsequent structure-function studies revealed that kinase null mutations as well as deletion of the putative TNFR domain complemented the *acr4-2* null mutant phenotype^18,20,24,25^. In contrast, partial deletion of the putative β-propeller domain or mutation of the conserved Cys180 in the β-propeller to tyrosine could not rescue the *acr4-2* phenotype^20,26^, suggesting an important functional role for the N-terminal segment of the ACR4 ectodomain.

β-propeller domains are often involved in protein – ligand or protein – protein interactions^27^ and thus different interaction partners for ACR4 ectodomain have been proposed, following the identification of root specific functions for ACR4^28,29^. Specifically, the plant peptide hormone CLAVATA3/ESR-RELATED 40 (CLE40) controls expression of the transcription factor WUSCHEL RELATED HOMEOBOX 5 (WOX5) to regulate root stem cell proliferation^29^. CLE40’s signaling capacity depends on the presence of ACR4 and ACR4 has been proposed to act as a direct receptor for CLE40 in the root^29–31^. Moreover, ACR4 has been reported to physically interact with the CLAVATA3 (CLV3) / CLE peptide receptor CLAVATA1 (CLV1), forming heteromeric complexes at the plasma-membrane^32^. In the same study, ACR4 homo-oligomers were observed^32^. PROTEIN PHOSPHATASE 2A-3 (PP2A-3) and WOX5 have been identified as direct interaction partners for the ACR4 cytoplasmic domain^33,34^. Here, we uncover the architecture of plant-unique CRINKLY4 RKs by solving crystal structures of ACR4 and *Physcomitrella patens*^35^ PpCR4.

## Results

For protein X-ray crystallographic analysis, we produced the ectodomains of ACR4 (ACR4^WD40-CRD^, residues 1 – 423) and PpCR4 (PpCR4^WD40-CRD^, residues 1 – 405), the isolated β-propeller domain of ACR4 (ACR4^WD40^, residues 1 – 334) and the kinase domain of ACR4 (ACR4^kinase^, residues 497 – 792) by secreted and cytoplasmic expression in insect cells, respectively. (see Methods) (Fig. 1a). All proteins were purified to homogeneity and the autophosphorylation activity of ACR4^kinase^ could be confirmed (Fig. 1b,c). No crystals were obtained for ACR4^kinase^ and initial crystals of ACR4^WD40-CRD^ and ACR4^WD40^ diffracted poorly. Enzymatic deglycosylation of ACR4^WD40^ yielded a new crystal form diffracting to 1.95 Å resolution. The structure was determined using the multiple anomalous dispersion method on a single crystal derivatized with a platinum compound (see Methods, Supplementary Table 1). Next, enzymatic deglycosylation of PpCR4^WD40-CRD^ yielded crystals diffracting to 2.7 Å resolution, enabling us to trace the entire CRINKLY4 ectodomain (Supplementary Table 1).

**Figure 1.**
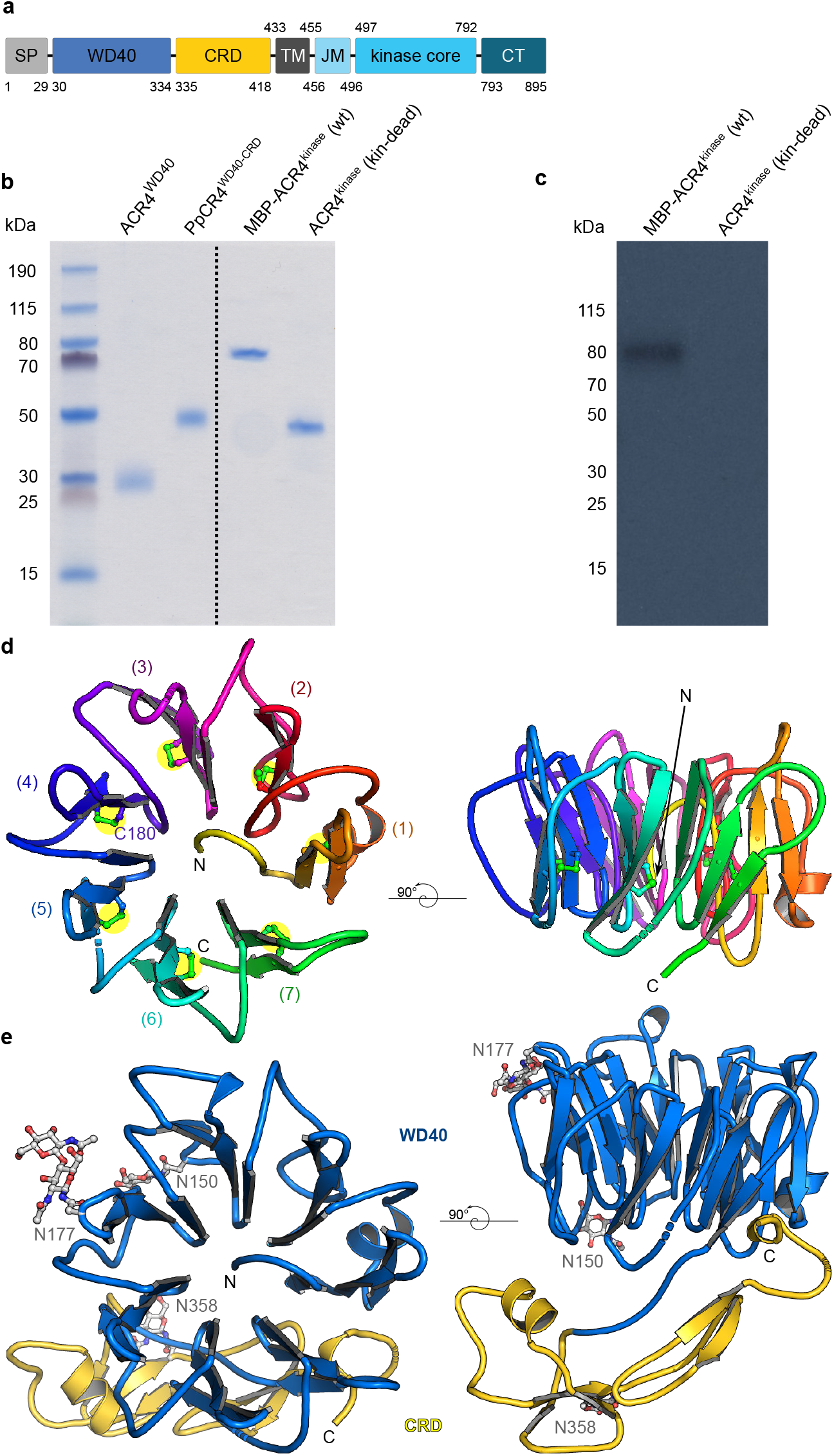
CRINKLY 4 receptor kinases harbor structurally unique β-propeller and cysteine-rich domains. **a**, ACR4 domain scheme: SP, signal peptide; WD40, WD40 domain; CRD, cysteine-rich domain; TM, transmembrane helix; JM, juxtamembrane region; CT, C-terminal tail. **b**, SDS-PAGE analysis of purified CRINKLY4 proteins expressed in insect cells. **c**, Autoradiography *in vitro* kinase assay of the wild-type ACR4 kinase domain fused to maltose-binding protein (MBP), and of the unfused kinase domain carrying a point mutation (Asp659→Asn) in the active site. The coomassie-stained gel loading control is shown in b (lanes on the right of the dotted line). **d**, Ribbon diagrams of ACR4^WD40^ in two orientations and colored from N-(yellow) to C-terminus (green). Disulfide bonds are shown in bonds representation and highlighted by yellow circles. **e**, Structure of PpCR4^WD40-CRD^ shown in two different orientations and colored in blue (WD40 domain) and yellow (CRD), respectively. The N-glycans visible in the electron density map are depicted in bonds representation (in gray).c

The N-terminal β-propeller domain of ACR4 and PpCR4 folded into a seven-bladed WD40 domain^27^ (Fig. 1d), as previously speculated^20^. Each blade is stabilized by a highly conserved disulfide bridge and connected by small loop regions, possibly an evolutionary adaptation to the extracellular environment (Fig. 1d, Supplementary Fig. 1). Cys180, which is found mutated to tyrosine in the *acr4-7* mutant^20^, forms a disulfide bond in the 4^th^ blade (Fig. 1d). The N- and C-terminal blades are not connected by disulfide bonds (Fig. 1d). The most N-terminal β-strand is buried in the center of the propeller and is highly conserved among all known CRINKLY4 receptors^36^ (Fig. 1d, Supplementary Fig. 1). Several small loops connecting the different blades of the WD40 domain appear partially disordered in our ACR4 and PpCR4 structures (Fig. 1d,e).

The C-terminal CRD comprises PpCR4 residues 313-401 and folds into a well defined β-sandwich structure stabilized by six invariant disulfide bridges (Fig. 1e, Supplementary Fig. 1, see below). The WD40 and CRD domains are connected by a short linker region (Fig. 1e). Analysis of crystal lattice arrangements with the program PISA^37^ and analytical size-exclusion chromatography experiments (Supplementary Fig. 2) together indicate that the ACR4 and PpCR4 ectodomains behave as monomers in solution. All surface exposed cysteines in ACR4 and PpCR4 contribute to disulfide bond formation (Fig. 1d,e; Supplementary Fig. 1). The N-glycosylation pattern differs between ACR4 and PpCR4 (Fig. 1e, Supplementary Fig. 1). Taken together, a compact WD40 and a cysteine-rich domain represent structural fingerprints of monomeric CRINKLY4 ectodomains.

Structural homology searches against ACR4^WD40^ using the program DALI^38^ returned the extracellular WD40 domain of the secreted β-lactamase inhibitor protein II BLIP-II from the soil bacterium *Streptomyces exfoliatus* as top hit (DALI Z-score 23.2, root mean square deviation [r.m.s.d.] is ∼2.2 Å comparing 192 corresponding C_α_ atoms) (Supplementary Fig. 3)^39^. A previously reported homology model of ACR4^WD40^ had been based on the BLIP-II structure^20^. The UV-B photoreceptor UV-B – RESISTANCE 8 (UVR8) represents the closest structural homolog in plants (Dali Z-score 22.1, r.m.s.d. is ∼2.4 Å comparing 218 corresponding C_α_ atoms) (Supplementary Fig.3)^40^. ACR4^WD40^ however lacks the UVR8 tryptophan cage involved in UV-B light sensing^40,41^ and both BLIP-II and UVR8 are devoid of the buried N-terminal strand and the conserved disulfide bridge pattern present in ACR4^WD40^. Thus, the pore-filling N-terminus and the invariant blade disulfide bonds are unique structural features of extracellular CRINKLY4 WD40 domains.

We next studied the interaction of ACR4^WD40-CRD^ with its proposed ligand CLE40^29–32^. As ACR4 has been previously reported to form hetero-oligomers with the LRR-RK CLV1, we sought to include the CLV1 ectodomain in these experiments, but we could not produce well-behaving protein samples of the AtCLV1 ectodomain by secreted expression in insect cells (Supplementary Fig. 4), and consequently could not use the CLV1 ectodomain for biochemical or crystallographic experiments. We thus replaced CLV1 with the LRR ectodomain of the sequence-related CLE peptide receptor BARELY ANY MERISTEM (BAM1) in our *in vitro* binding experiments (Fig. 2a). We found that CLE40 binds the AtBAM1 ectodomain with a dissociation constant (K_*d*_) of ∼1 μM (Fig. 2b) but shows no detectable binding to the ACR4 ectodomain in quantitative grating-coupled interferometry (Fig. 2b) and isothermal titration calorimetry (Fig. 2c) assays. Thus, CLE40 does not represent a direct ligand for the ACR4 ectodomain.

**Figure 2.**
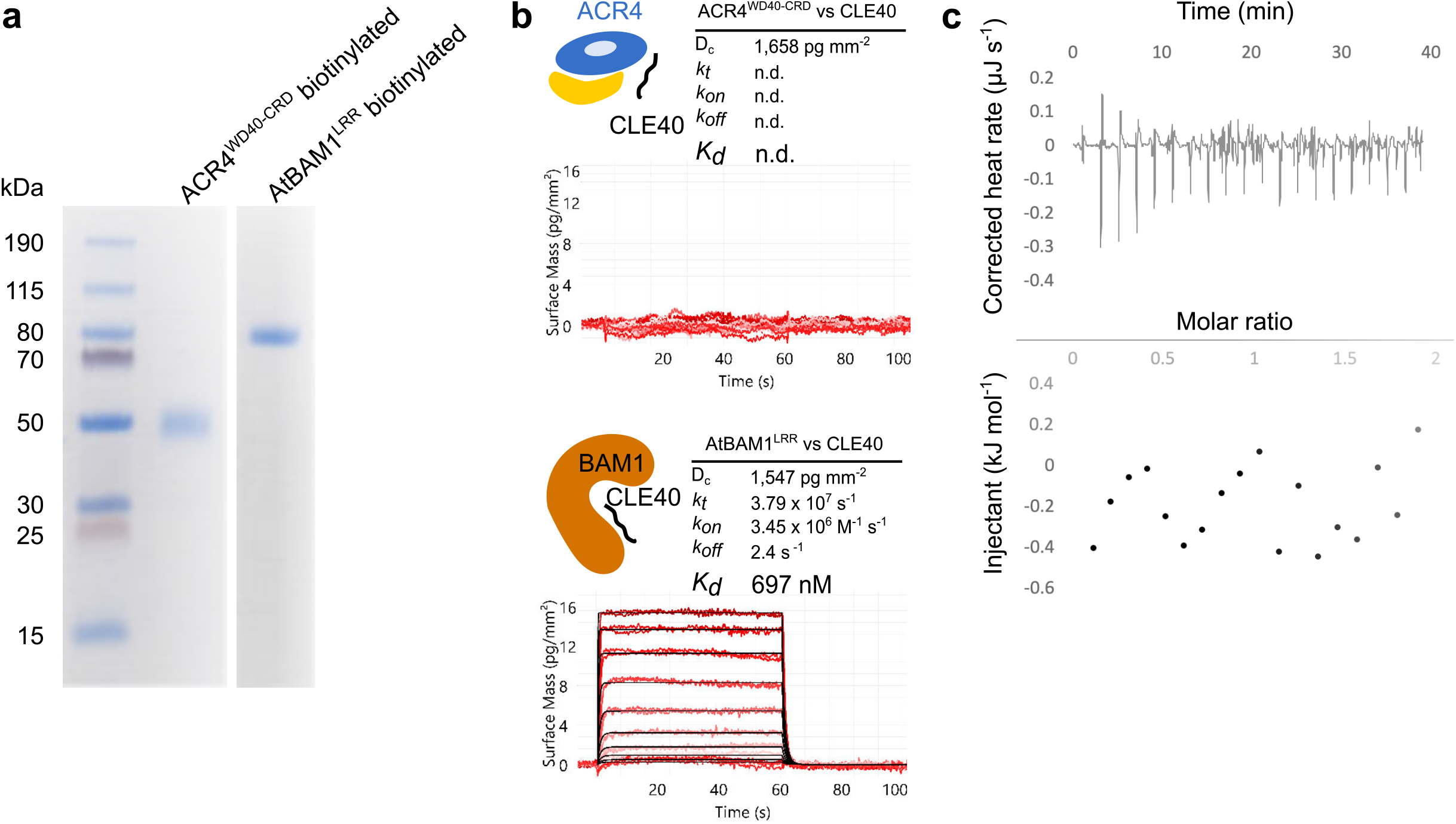
The ACR4 ectodomain does not bind the peptide hormone CLE40 *in vitro*. **a**, SDS-PAGE analysis of the biotinylated ACR4^WD40-CRD^ and AtBAM1^LRR^ ectodomains used for binding experiments. **b**, Quantitative grating-coupled interferometry (GCI) binding assay of a synthetic CLE40 peptide versus ACR4^WD-CRD^ and BAM1^LRR^. Shown are sensorgrams with raw data in red and their respective fits in black. Table summaries of kinetic parameters are shown alongside (D_c_, density of captured protein; k_t_, mass transport coefficient; k_on_, association rate constant; k_off_, dissociation rate constant; K_d_, dissociation constant; n.d., no detectable binding, n=3). **c**, Isothermal titration calorimetry (ITC) experiment of ACR4^WD-CRD^ versus CLE40. No binding was detected in this assay (n=3).

Using the previously documented seed retardation phenotype of the *acr4-2* mutant^18,20^ we next carried out genetic complementation analyses using different constructs expressed from the *ACR4* promoter. In agreement with an earlier report^20^, a construct in which the entire cytoplasmic domain of ACR4 had been deleted could not rescue the seed development phenotypes of *acr4-2* plants (Fig. 3a). Full-length ACR4 lacking kinase activity partially restored seed development in *acr4-2* plants (Fig. 3a). Strikingly, expression of full-length PpCR4, the ectodomain of which shares only 40% sequence identity at the amino-acid level with ACR4^WD40-CRD^, from the ACR4 promoter could partially complement *acr4-2* phenotypes as well. Together, these experiments reinforce an evolutionary conserved function for CRINKLY4 RKs, which are however not strictly dependent on the protein kinase activity of the receptor.

**Figure 3.**
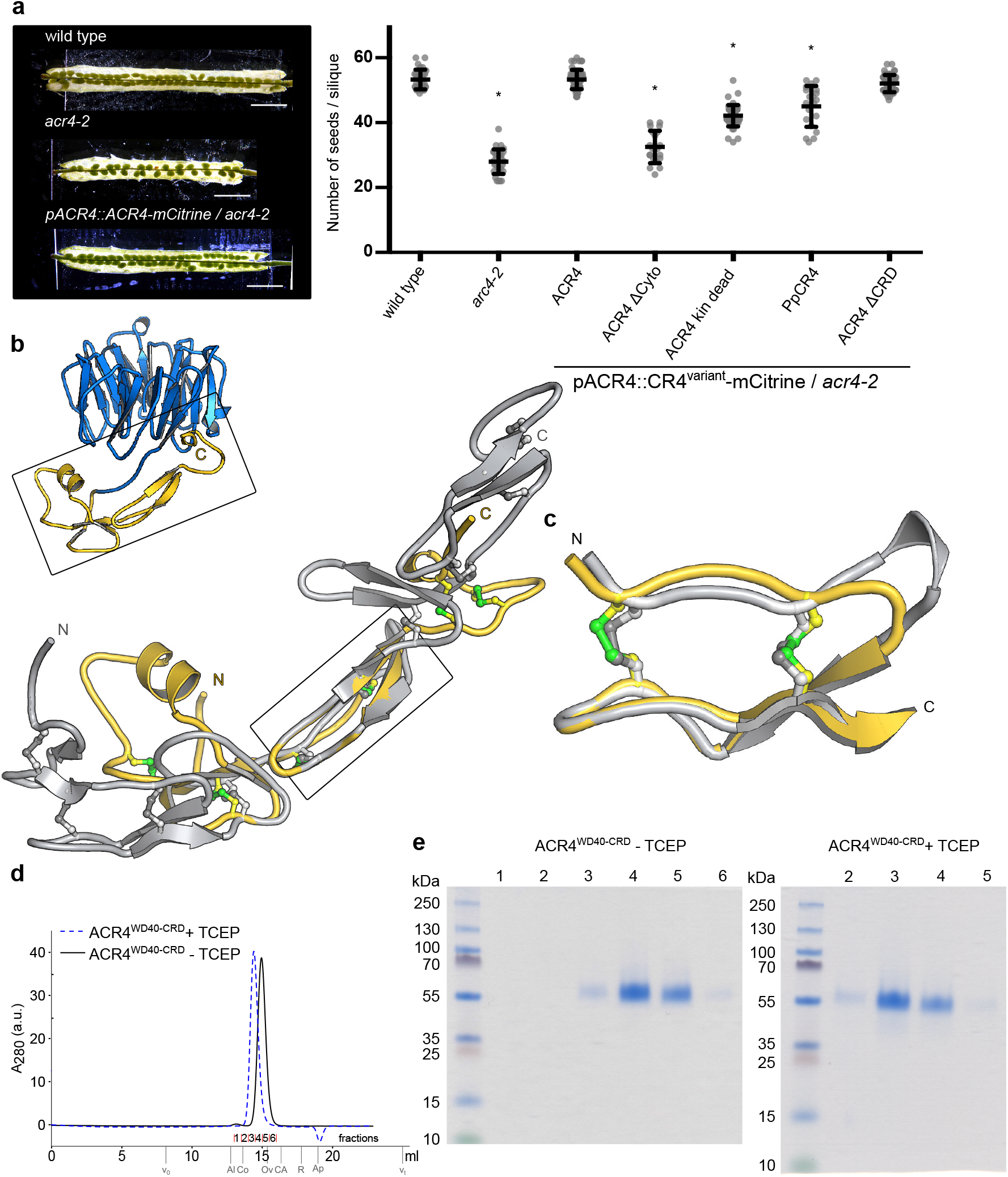
CRINKLY4 ectodomains harbor an evolutionary conserved function. **a**, Reverse genetic rescue experiments of the seed development phenotype of *acr4-2*. Left panel: Seed development phenotypes of wild type, *acr4-2* and a complemented line. Right panel: Ten siliques per transgenic line from three independent homozygous T3 lines were pooled and plotted as beeswarm plots with the bold line representing mean, whiskers indicating the standard deviation, and circles depicting the raw data. Seed counts per silique significantly different from wild type were determined by simultaneous comparisons of several mutants against wild type using the Dunnett procedure (indicated by an asterisk). **b**, Ribbon diagram overview of PpCR4^WD40-CRD^ (colors as in Fig. 1) and close-up view of the CRD superimposed to a type I TNF receptor ectodomain (PDB-ID 1NCF^77^; in gray). The six invariant disulfide bridges of CRINKLY4 CRDs are shown in green, the disulfide bonds in TNFR are shown in gray (in bonds representation). **c**, Superposition of the structurally homologous PpCR4^CRD^ (in yellow) and TNFR (in gray) core segments (r.m.s.d. is ∼1 Å comparing 20 corresponding C_α_ atoms). **d**, Analytical size-exclusion chromatography of ACR4^WD40-CRD^ in the pre- or absence of Tris(2-carboxyethyl)phosphine (TCEP). Void (V_0_), total (V_t_), and elution volumes for molecular-mass standards (Al, Aldolase, 158 kDa; Co, Conalbumin, 75 kDa; Ov, Ovalbumin, 44 kDa; CA, Carbonic Anhydrase, 29 kDa; R, Ribonuclease A, 13.7 kDa; Ap, Aprotinin; 6.5 kDa) are indicated. **e**, SDS-PAGE analysis of fractions shown in d.

The 2.7 Å crystal structure of the entire ectodomain from PpCR4 enabled us to further characterize the ∼90 amino-acid CRINKLY4 CRD (Fig. 3b). A structural homology search with DALI^38^ indeed identified several TNFR domains as top hits, but with very low DALI Z-scores (4.1-2.9). Structural superposition of PpCR4^CRD^ with the previously reported structure of a type I TNF receptor extracellular domain revealed that only a small portion of the CRINKLY4 aligns with canonical TNFR domains (r.m.s.d. is ∼1 Å comparing 20 corresponding C_α_ atoms, Fig. 3b). The segment includes a small β-hairpin and two conserved disulfide bridges located at the center of the CRINKLY4 CRD (Fig. 3c). Structural superposition of the eight molecules in the asymmetric unit of our PpCR4^WD40-CRD^ crystal structure (Supplementary Table 1) revealed only subtle movements of the CRD versus the WD40 domain (r.m.s.d. is ∼0.3-0.5 Å comparing 360 corresponding C_α_ atoms, Supplementary Fig. 5). In line with this, we located a small WD40 – CRD domain interface using PISA^37^ (total buried surface area is ∼900 Å^2^). The interface is formed by the C-terminus of the CRD (PpCR4 residues 385-401) that makes mainly hydrophobic interactions with a small groove located between the N- and C-terminal blade of the WD40 domain (Supplementary Fig. 6). Additional contacts originate from a small α-helix in the CRD and several loop regions in PpCR4^WD40^ (Supplementary Fig. 6).

Using the now experimentally determined domain boundaries of the ACR4 CRD (Supplementary Figs. 7 and 1), we re-performed complementation assays of the *acr4-2* mutant with a construct in which the entire CRD was omitted (ACR4 ΔCRD). As previously reported^20,24^, we found that ACR4 ΔCRD can rescue the seed development phenotype of *acr4-2* plants (Fig. 3a). Recently, mutation of the cysteine residues in ACR4^WD40^ and ACR4^CRD^ involved in the formation of disulfide bonds in our structures (Fig.1, 3b,c) to alanine resulted in a functional receptor for seed development^24^. We monitored migration of the purified ACR4^WD40-CRD^ ectodomain under oxidizing and strongly reducing conditions in analytical size exclusion chromatography experiments and found that reduction of ACR4^WD40-CRD^ did not induce aggregation of the receptor (Fig. 3d,e). Together, the CRINKLY4 CRD only shares weak structural homology with animal TNFR domains, has a conserved domain interface with the WD40 domain and is dispensable for seed development. The conserved disulfide bonds appear to be involved in structural stabilization. The domain interface between the WD40 domain and the CRD is conserved among CRINKLY4 receptors from different species (Supplementary Figs. 1, 6).

While the CRD domain appears to be dispensable for at least some of ACR4’s physiological functions, our and previous findings^20,24^ argue for an important role of the structurally unique WD40 domain in CRINLKY4 receptors. We located evolutionary conserved, surface exposed residues at the ‘back side’ of the ACR4 WD40 domain (Fig. 4a, Supplementary Fig. 1), which in our PpCR4^WD40-CRD^ structure is in contact with the CRD (Fig. 1e). We replaced individual residues by alanine or glutamine, respectively and assessed the ability of the resulting mutant proteins to complement the *acr4-2* seed development phenotype (Fig. 4a-c). We analyzed three independent homozygous T3 lines per mutant receptor and found that most mutations behaved similar to wild type (Fig. 4b) and that none of mutants tested displayed the strong loss-of-function phenotype of *acr4-2* plants (Fig. 4b,c). Plants in which either Tyr157 or Asn158/Asn196 were mutated had seed numbers per silique that were significantly reduced compared to wild type (Fig. 4b). While there was no electron density for a N-glycan at position Asn158 in the ACR4^WD40^ structure (see Methods), the corresponding Asn150 in PpCR4 was found glycosylated (Fig. 1e). ACR4 Asn196 is predicted to be N-glycosylated as well^42^, suggesting that the weak loss-of-function phenotypes observed in our Tyr157/Asn158 and Asn196 point mutants may be caused by an altered N-glycosylation pattern of the receptor (Fig. 4b,c).

**Figure 4.**
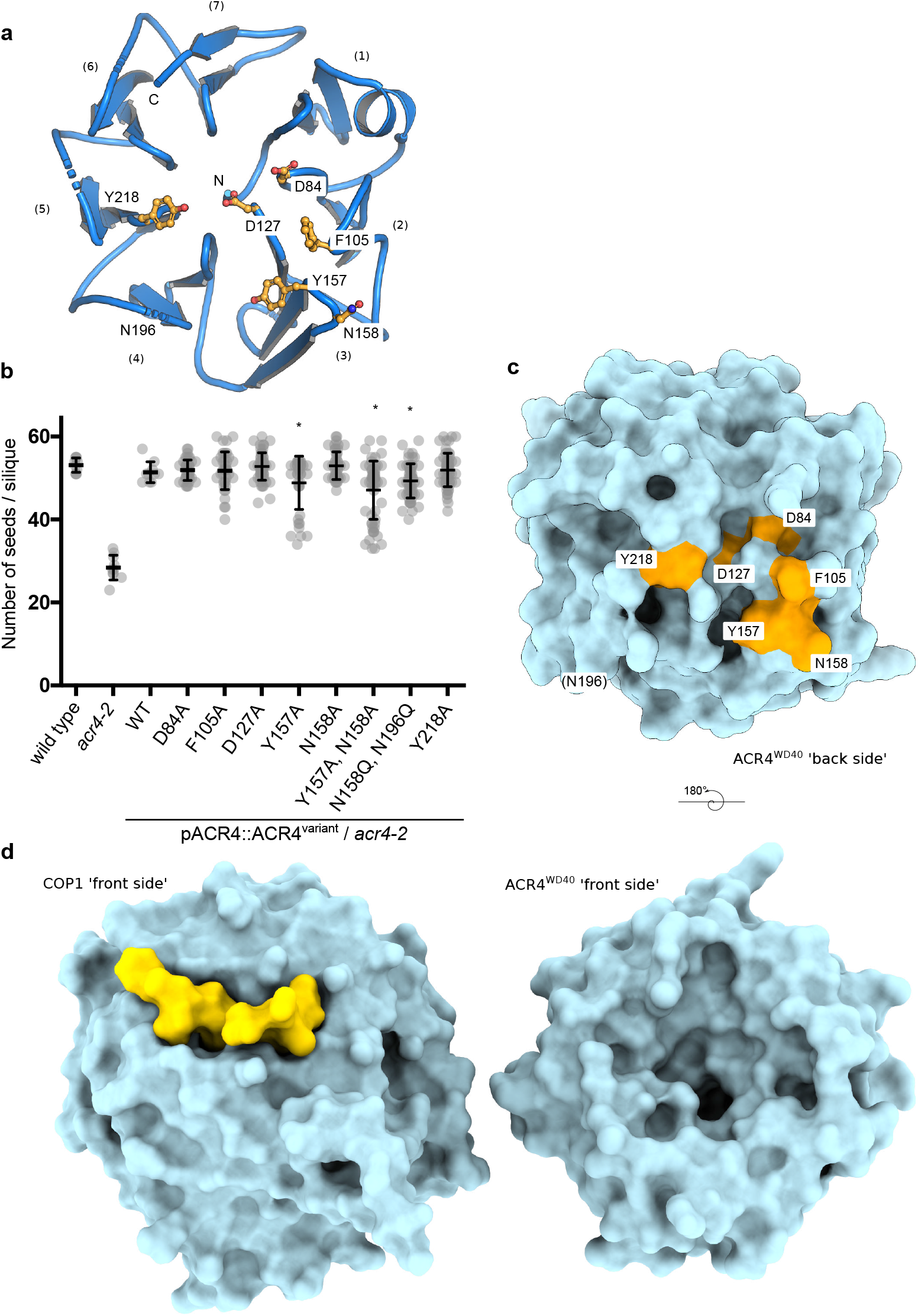
The CRINKLY4 WD40 domain contains a putative ligand binding groove. **a**, Ribbon diagram of ACR4^WD40^ (in blue) with surface exposed conserved residues shown in bonds representation (in orange) at the exposed surface. Blade numbers are indicated. **b**, Effect on surface point-mutations on ACR4-mediated seed production. Ten siliques per transgenic line from three independent homozygous T3 complementation lines were pooled and plotted as beeswarm plots with the bold line representing mean, whiskers indicating the standard deviation, and circles depicting the raw data. The plots for wild type, *acr4-2* and *ACR4* were generated from same data sets shown in Fig. 3a. Seed counts per silique significantly different from wild type were determined by simultaneous comparisons of several mutants against wild type using the Dunnett procedure (indicated by an asterisk). **c**, Molecular surface of the ACR4^WD40^ β-propeller domain ‘back side’ (in light blue). The positions of the mutated residues are highlighted in orange. **d**, Comparison of the ‘front sides’ of the structurally related WD40 domains of COP1 (PDB-ID 6QTO^43^ left panel) and ACR4 (right panel, r.m.s.d is ∼3.5 comparing 205 corresponding C_α_ atoms). The COP1 VP-peptide ligand derived from the transcription factor HY5 is shown in yellow. Note the large and deep putative binding groove in the corresponding surface area in ACR4^WD40^.

We next analyzed the molecular surface of the ‘front side’ of ACR4^WD40^, which represents another canonical binding surface for peptide and protein ligands in many cytoplasmic or nuclear localized WD40 proteins^27^. We located a large binding groove in ACR4^WD40^ formed by the WD40 domain core and by small surrounding loop regions, which appear similar in our ACR4^WD40^ and PpCR4^WD40-CRD^ WD40 domain structures (r.m.s.d. is ∼1.4 Å comparing 246 corresponding C_α_ atoms, Supplementary Fig. 7). The very low degree of sequence surface conservation in the putative binding groove in apo ACR4^WD40^ renders mutational analysis of the full-length receptor *in planta* difficult (Supplementary Figs. 1, 8). The binding groove is however larger and deeper compared to the VP-peptide binding site in the structurally related WD40 domain of the light-signaling E3 ubiquitin ligase CONSTITUTIVELY PHOTOMORPHOGENIC 1 (COP1) (Fig. 4d)^43^. It may thus provide and interaction platform for high molecular weight ligand.

## Discussion

Our crystal structures (Fig. 1d,e) and reverse genetic analyses (Fig. 3a) together reveal an evolutionary conserved domain architecture for plant-unique CRINKLY4 receptor kinases^36^. The CRINKLY4 WD40 domain differs from known cytoplasmic and extracellular WD40 domains^27,39,40,43^, with its seven blades being stabilized by disulfide bridges and the hydrophobic core of the domain being reinforced by insertion of the protein’s N-terminus (Fig. 1d,e). We speculate that these unique structural features represent an adaptation to CRINKLY4 ectodomains being exposed to the plant cell wall environment. Previous^20^ and our genetic data argue for an important function for the ACR4 WD40 domain in seed development (Fig. 3a). A large groove located on the ‘front side’ of ACR4 may be involved in the binding of a ligand (Fig. 4d). This ligand could be a small molecule, a protein or a peptide, and may be larger than the octameric peptide motifs recognized by COP1 (Fig. 4d). The low degree of sequence conservation of residues contributing to the formation of the binding groove in the WD40 domain (Fig. 4d, Supplementary Fig. 1) and the fact that PpCR4 can functionally replace ACR4’s function in Arabidopsis seed development (Fig. 3a) together indicate that CRINKLY4 receptors may sense a family of structurally conserved ligands.

Our quantitative binding assays reveal that the previously proposed peptide ligand CLE40 cannot directly interact with the ACR4 ectodomain (Fig. 2), but we cannot rule out that CLE40 binds the CLV1 ectodomain in a signaling complex also containing ACR4^29,30,32^. The architecture and cellular functions of CLV1 – ACR4 signaling complexes remain to be elucidated, with recombinant expression and purification of the CLV1 ectodomain representing a significant challenge (Supplementary Fig. 4). BAM1 cannot fully replace CLV1 in quantitative biochemical assays, as it binds CLE40 only with moderate affinity (Fig. 2). In contrast, the CLE family member CLE9 binds BAM1 with nanomolar affinity^44,45^. In solution and in the absence of ligand, CRINKLY4 ectodomains behave as monomers (Fig. 3e, Supplementary Fig. 2). The previously observed ACR4 homo-oliogomers^32^ may thus be generated by ligand-induced oligomerisation of several CRINKLY4 ectodomains and/or be stabilized by interaction of the CRINKLY4 transmembrane helices, as previously suggested^46,47^.

Analysis of the CRINKLY4 cysteine-rich domain revealed only weak structural homology with animal TNFR domains (Fig. 3b)^14^. In line with this, we could not locate proteins with homology to tumor necrosis factors in the *Arabidopsis* or *Physcomitrella patens* genomes^48,49^. The CRINKLY4 CRD contains six conserved disulfide bridges (Fig. 1e, Supplementary Fig. 1), which in our PpCR4^WD40-CRD^ structure appear to be involved in structural stabilization (Fig. 3b,c). However, CRINKLY4 ectodomains can withstand reducing conditions (Fig. 3e), and thus the putative function of the CRD could indeed be regulated by changes in the cell wall redox environment^24^.

Enzymatic assays of the CRINKLY4 cytoplasmic domains obtained from prokaryotic^18,21–23^ or eukaryotic expression hosts (Fig. 1b,c) clearly identify CRINKLY4s as active protein kinases. Our and previous^20^ reverse genetic experiments suggest that the ACR4 cytoplasmic domain has to be present for normal seed development in Arabidopsis, yet its catalytic activity seems to be dispensable (Fig 1e). Similar observations have been made for CrRLK1L-family receptor kinases^50–52^. The mechanistic implications are poorly understood, but the involvement of protein phosphatases in both CR4 and CrRLK1L-mediated signal transduction^33,53,54^ suggests that the cytoplasmic kinases domains of these receptors may act as scaffolding proteins that can become phosphorylated despite not requiring auto- and trans-phosphorylation activity themselves.

Genetic interactions between ACR4 and other receptor kinases such as ABNORMAL LEAF SHAPE 2 (ALE2)^55^ the LRR-RKs CLV1^29,30,32^ and GSO1/GSO2^45,56^ have so far not yielded a mechanistic understanding of CRINKLY4’s signaling functions. Also, no ligand candidate for ACR4 or for its homologs in Arabidopsis has emerged from forward genetic screens^21^. Our identification of a putative ligand binding pocket in ACR4^WD40^ now reinforces the notion that *bona fide* ligands for CR4s may exist and that their identification may be achieved using a combination of genetic and biochemical approaches.

## Material and Methods

### Protein expression and purification

ACR4 coding sequences for the WD40 domain (residues 1 – 334) and its entire ectodomain (residues 1 – 423) were amplified from *A. thaliana* cDNA. PpCR4^WD40-CRD^ (residues 1 – 405), AtCLV1 (residues 25 – 621) and BAM1 (residues 20 – 637) were synthesized by Geneart (Germany) with codon usage optimized for expression in *Trichoplusia ni*. The constructs of ACR4 and PpCR4 were cloned in a modified pFastBac vector (Geneva Biotech), containing a TEV (tobacco etch virus protease) cleavable C-terminal StrepII – 9x His tag. ACR4^WD40-CRD^, CLV1 and BAM1 were also cloned into the vector holding a native signal peptide or the *Drosophila melanogaster* BiP secretion signal peptide, respectively, a C-terminal TEV cleavable StrepII – 10x His tag and a non-cleavable Avi-tag^57,58^. *Trichoplusia ni* (strain Tnao38) cells^59^ were infected with a multiplicity of infection (MOI) of 1 at a density of 2 x 10^6^ cells ml^-1^ and incubated 26h at 28 °C and 48h at 22 °C. The secreted protein was purified from the supernatant by Ni^2+^ affinity chromatography on a HisTrap Excel column (GE healthcare), equilibrated in 50 mM KP_i_ pH 7.6, 250 mM NaCl, 1 mM 2-Mercaptoethanol, followed by StrepII affinity chromatography on a Strep-Tactin XT Superflow high affinity column (IBA), equilibrated in 20 mM Tris pH 8.0, 250 mM NaCl, 1 mM EDTA. The tag was cleaved with His-tagged TEV protease at 4 °C overnight and removed by a second Ni^2+^ affinity chromatography step. Proteins were then further purified by size-exclusion chromatography on either a Superdex 200 increase 10/300 GL, Hi Load 16/600 Superdex 200 pg, or a HiLoad 26/600 pg column (GE Healthcare), equilibrated in 20 mM HEPES pH 7.5, 150 mM NaCl. For crystallization, ACR4^WD40^ and PpCR4^WD40-CRD^ were dialyzed in 20 mM sodium citrate pH 5.0, 150 mM NaCl and treated with Endoglycosidase H, F1, and F3 to cleave sugar chains. Proteins were then purified by ion exchange chromatography on a HiTrapSP HP column (GE Healthcare), equilibrated in 20 mM Citrate pH 5.0, 25 mM NaCl for ACR4^WD40^ or 20 mM Citrate pH 3.5, 25 mM NaCl for PpCR4^WD40-CRD^, respectively. Fractions were pooled, concentrated and further purified by size-exclusion chromatography.

### *In vitro* kinase phosphorylation assay

Coding sequence of ACR4 kinase domain (residues 497 – 792) was amplified from *A. thaliana* cDNA and cloned in a mofidied pFastBac vector harboring a TEV cleavable N-terminal maltose binding protein (MBP) – StrepII – 10x His tag. Point mutation was introduced into the ACR4 (Asp659→Asn; hereafter ACR4^D659N^, Supplementary Table 2) coding sequence using the primer extension method for site-directed mutagenesis, rendering the kinase inactive^60^. Insect cells were infected with a MOI of 1 at a density of 2 ⨯ 10^6^ cells ml^-1^ and incubated 26h at 28 °C and 48h at 22 °C. Cells were pelleted by centrifugation at 4,000 x g, 4 °C for 15 min and resuspended in buffer A (20 mM HEPES pH 7.5, 500 mM NaCl, 4 mM MgCl_2_ and 2 mM 2-Mercaptoethanol) supplemented with 50 µg ml^-1^ DNAse I, 10 %(v/v) glycerol and 1 tablet of protease inhibitor cocktail (cOmplete, Roche), followed by sonication. The cell lysate was centrifuged at 35,000 x g, 4 °C for 60 min and the protein was purified from the supernatant by Ni^2+^ affinity chromatography with buffer A, followed by StrepII affinity chromatography. For ACR4^D659N^, the 10x His – StrepII – MBP tag was cleaved with His-tagged TEV protease at 4 °C overnight and removed by Ni^2+^ affinity chromatography. Proteins were then further purified by size-exclusion chromatography on a Superdex 200 increase 10/300 GL column equilibrated in 20 mM Tris-HCl pH 8, 250 mM NaCl, 4 mM MgCl_2_ and 0.5 mM TCEP. Monomeric peak fractions were collected and concentrated for analyses. For *in vitro* kinase assays, 2 µg of MBP-ACR4 and 1 µg of ACR4^D659N^ were used in a reaction volume of 20 µl. The reactions were started by addition of 5 µCi [γ-^32^P]-ATP (Perkin-Elmer, Waltham, MA), incubated at room temperature for 45 min and terminated by the addition of 6x SDS loading dye, immediately followed by heating the samples at 95 °C. Proteins were separated by SDS-PAGE in 4 – 15 % gradient gels (TGX, Biorad) and ^32^P-derived signals were visualized by exposing the gel to an X-ray film (SuperRX, Fujifilm).

### Crystallization and data collection

Crystals of the deglycosylated ACR4^WD40^ and PpCR4^WD40-CRD^ developed at room temperature in hanging drops composed of 1 µl protein solution (ACR4^WD40^, 20 mg/ml; PpCR4^WD40-CRD^, 16 mg/ml) 1 µl of crystallization buffer (16 % PEG 6,000, 0.01 M tri-sodium citrate pH 5.0 for ACR4^WD40^; 15 % PEG 4,000, 0.2 M imidazole malate pH 7.0 in the case of PpCR4^WD40-CRD^) suspended above 1.0 ml of the latter as reservoir solution and using microseeding protocols. Crystals were cryo-protected by serial transfer into crystallization buffer supplemented with 20 % (v/v) ethylene glycol and snap-frozen in liquid nitrogen. For heavy-atom derivatization, crystals of ACR4^WD40^ were transferred in the crystallization buffer containing 2 mM K_2_[Pt(CNS)_6_] and incubated for 2.5h. Crystals were cryo-protected by serial transfer into crystallization buffer supplemented with 20 % (v/v) glycerol and cryo-cooled in liquid nitrogen. Platinum multi-wavelength anomalous diffraction (MAD) data were collected to 3.2 Å resolution was collected at beam-line PXIII at the Swiss Light Source (SLS), Villigen, CH. A native data for ACR4 ^WD40^ and PpCR4^WD40-CRD^ were recorded at a resolution of 1.95 Å and 2.70 Å, respectively (Supplementary Table 1). Data processing and scaling were done with XDS and XSCALE^61^.

### Structure solution and refinement

Nine consistent Pt sites were located in three wavelength MAD data using the program SHELXD^62^ followed by site refinement and phasing in SHARP^63^. The resulting heavy atom sites and starting phases (FOM was 0.35 to 3.2 Å resolution) were input into phenix.autobuild^64^ for non-crystallographic symmetry (NCS) averaging, phase extension, density modification (FOM was 0.75 to 1.95 Å resolution) and iterative model building. The refined (Refmac5^65^) model comprises four ACR4^WD40^ molecules in the asymmetric unit with an associated solvent content of 0.42. The space group *P* 2_1_ with a β angle of 90.1° was validated using the programs POINTLESS^66^ and ZANUDA^67^. The structure of PpCR4^WD40-CRD^ was solved using the molecular replacement method using an ACR4^WD40^ monomer as search model in calculations with the program PHASER^68^. The solution comprises eight PpCR4^WD40-CRD^ molecules in the asymmetric unit. The structure was completed in alternating cycles of manual model building in COOT^69^ and restrained NCS refinement in phenix.refine^70^. Ile156 represents a Ramachandran plot outlier in each chain, but is well defined by electron density. Structural diagram were prepared in Pymol (https://sourceforge.net/projects/pymol/) and ChimeraX^71^.

### Biotinylation of proteins

The respective proteins (20 – 100 µM) were biotinylated with biotin ligase BirA^58^ (2 µM) for 1h at 25 °C, in a volume of 200 µl; 25 mM Tris pH 8, 150 mM NaCl, 5 mM MgCl2, 2 mM 2-Mercaptoethanol, 0.15 mM Biotin, 2 mM ATP, followed by size-exclusion chromatography to purify the biotinylated proteins.

### Grating – coupled interferometry

GCI experiments were performed with the Creoptix WAVE system (Creoptix AG, Switzerland), using 4PCP WAVE chips (thin quasiplanar polycarboxylate surface; Creoptix, Switzerland). Chips were conditioned with borate buffer (100 mM sodium borate pH 9.0, 1 M NaCl; Xantec, Germany) and activated with 1:1 mix of 400 mM *N*-(3-dimethylaminopropyl)-*N’*-ethylcarbodiimide hydrochloride and 100 mM *N*-hydroxysuccinimide (Xantec, Germany) for 7 min. Streptavidin (50 µg ml^-1^; Sigma, Germany) in 10 mM sodium acetate pH 5.0 (Sigma, Germany) was immobilized on the chip surfaces and passivated with 0.5 % BSA (Roche, Switzerland) in 10 mM sodium acetate pH 5.0, followed by final quenching with 1M ethanolamine pH 8.0 (Xantec, Germany) for 7 min. Biotinylated ligands (20 – 50 µg ml^-1^) was captured by streptavidin immobilized on the chip surface. All kinetic analyses were performed at 25°C with a 1:2 dilution series from 10 µM of CLE40 peptides in 20 mM citrate pH 5.0, 250 mM NaCl, 0.01 % Tween 20. Blank injections were used for double referencing and a dimethylsulfoxide (DMSO) calibration curve for bulk correction. Analysis and correction of the obtained data was performed using the Creoptix WAVE control software (correction applied: X and Y offset; DMSO calibration; double referencing). Mass transport binding models with bulk correction were used. Experiments were performed in triplicates.

### Isothermal titration calorimetry

All ITC experiments were performed on a MicroCal PEAQ-ITC (Malvern Panalytical) with a 200 µl sample cell and a 40 µl injection syringe at 25 °C. Proteins were dialyzed into ITC buffer (20 mM sodium citrate pH 5.0, 250 mM NaCl) prior to all experiments. The CLE40 peptide (RQV[Hyp]TGSDPLHH) was synthesized (Peptide Specialty Labs GmbH) and dissolved directly in buffer. The dissolved peptide concentration was measured by right-angle light scattering (OMNISEC RESOLVE / REVEAL combined system, Malvern Panalytical). The protein concentrations were calculated based on their absorbance at 280 nm and their corresponding molar extinction coefficient. A typical experiment consisted of injecting 19 injections of 2 µl of 1000 µM CLE40 into the cell containing 100 µM ACR4. Experiments were performed in triplicates.

### Plant materials and generation of transgenic lines

*Arabidopsis thaliana* ecotype Columbia (Col-0) and SAIL_240_B04 (*acr4-2*^18^) were used for all experiments. *ACR4* gene (residues 1 – 895) and ACR4 promoter region (*pACR4*, 1847 bp upstream from ATG) were amplified from *A. thaliana* genomic DNA. *PpCR4* (residues 1 – 893) gene with *Physcomitrella patens* CDS was synthesized (Geneart, Germany). The coding sequences were cloned in a pDONR 221 Gateway vector (Invitrogen) and *pACR4* sequence was cloned in a pDONR P4-P1R Gateway vector (Invitrogen). *ACR4* variants carrying deletion or point mutations were generated using the primer extension method. pDONR P2R-P3 Gateway vector harboring mCitrine or 6x HA tag were used to attach C-terminal tag. Expression constructs were generated with LR Gateway Cloning (Invitrogen) in pH7m34GW^72^; pACR4::ACR4 (residues 1 – 895)-mCitrine, pACR4::ACR4_ΔCyto (residues 1 – 492)-mCitrine, pACR4::PpCR4 (residues 1 – 893)-mCitrine, pACR4::ACR4_ΔCRD (residues 1 – 895 with deletion 335 – 423)-mCitrine, pACR4::ACR4_K540R-HA, pACR4::ACR4_D84A-mCitrine, pACR4::ACR4_F105A-mCitrine, pACR4::ACR4_D127A-mCitrine, pACR4::ACR4_Y157A-mCitrine, pACR4::ACR4_N158A-mCitrine, pACR4::ACR4_Y157A, N158A-mCitrine, pACR4::ACR4_Y218A-mCitrine, pACR4::ACR4_N158Q, N196Q-HA. They were transformed in *acr4-2* backgound by floral dipping method^73^ with *Agrobacterium tumefaciens* strain GV3101 (Supplementary Table 3).

### Seed counting and statistical analysis

Plants were germinated on 0.5 MS (Murashige and Skoog) agar plates after 3 days in dark at 4°C. Seedling were transferred to soil and grown at 22°C, under long days (16 h light / 8 h dark) for 6 weeks. A top opened flower was defined as position 1 and a silique at position 12 was collected for analyses in a blind manner. 10 siliques were sampled for independent lines as biological replicates and seeds were counted under a stereo microscope. The simultaneous comparisons of the different transgenic lines vs wild type were performed using the Dunnett procedure^74^ for the primary endpoint number seeds per silique using the count transformation model^75^. The Comprehensive R Archive Network packages multcomp^76^ and cotram^75^ were used in R, version 3.6.3.

## Data availability

Data supporting the findings of this manuscript are available from the corresponding authors upon reasonable request. A reporting summary for this article is available as a Supplementary Information file. Coordinates and structure factors have been deposited in the Protein Data Bank (PDB) with accession codes 7A0J (ACR4^WD40^) and 7A0K (PpCR4^WD40-CRD^).

## Acknowledgments

We thank K. Lau for help with crystallographic data collection and U. Hohmann, D. Couto and J. Santiago for providing feedback on the manuscript. This work was supported by grants 31003A_176237 and 31CP30_180213 from the Swiss National Science Foundation (to MH), by a Howard Hughes Medical Institute (HHMI) International Research Scholar Award (to MH) and by a Human Frontier Science Program (HFSP) postdoctoral fellowship (to SO).

## Author contributions

MH and SO designed the study, SO performed all biochemical and genetic experiments, SO and MH phased and refined the structures, LAH performed the statistical analysis and SO and MH wrote the manuscript.

---

**Supplementary Figure 1.**
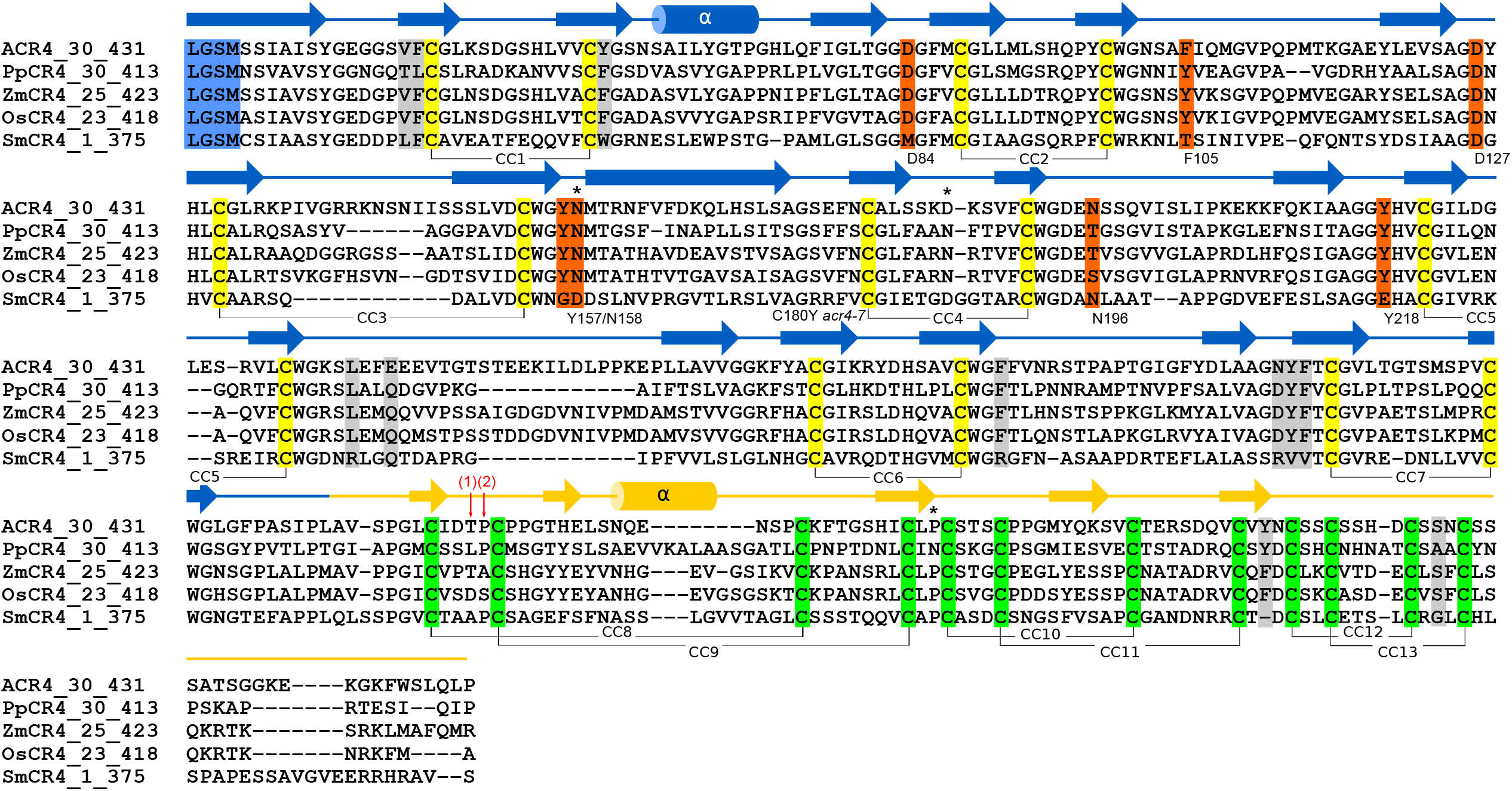
Structure-based multiple sequence alignment of CRINKLY4 receptor ectodomains from different species. Structure based T-COFFEE^78^ sequence alignment and including a secondary structure assignment calculated with DSSP^79^ (WD40 domain in blue, CRD in yellow). Invariant cysteine residues contributing to disulfide bonds in the WD40 domain or CRD domain are highlighted in yellow and green, respectively. Residues analyzed with point mutations in this study are shown in orange. Conserved residues in the WD40 – CRD domain interface are depicted in gray. Asterisks denote the location of experimentally confirmed N-glycosylation sites. Red arrows represent domain boundaries for the ΔTNFR/CRD deletion constructs in previous reports: (1)^24^, (2)^20^. ACR4 (*Arabidopsis thaliana*) UNIPROT-ID (http://uniprot.org) Q9LX29; PpCR4 (*Physcomitrella patens*) A9RKG8; ZmCR4 (*Zea mays*) O24585; OsCR4 (Oryza sativa) Q75J39; SmCR4 (*Selaginella moellendorffii*) D8T625. Note that the annotated SmCR4 sequence may be incomplete.

**Supplementary Figure 2.**
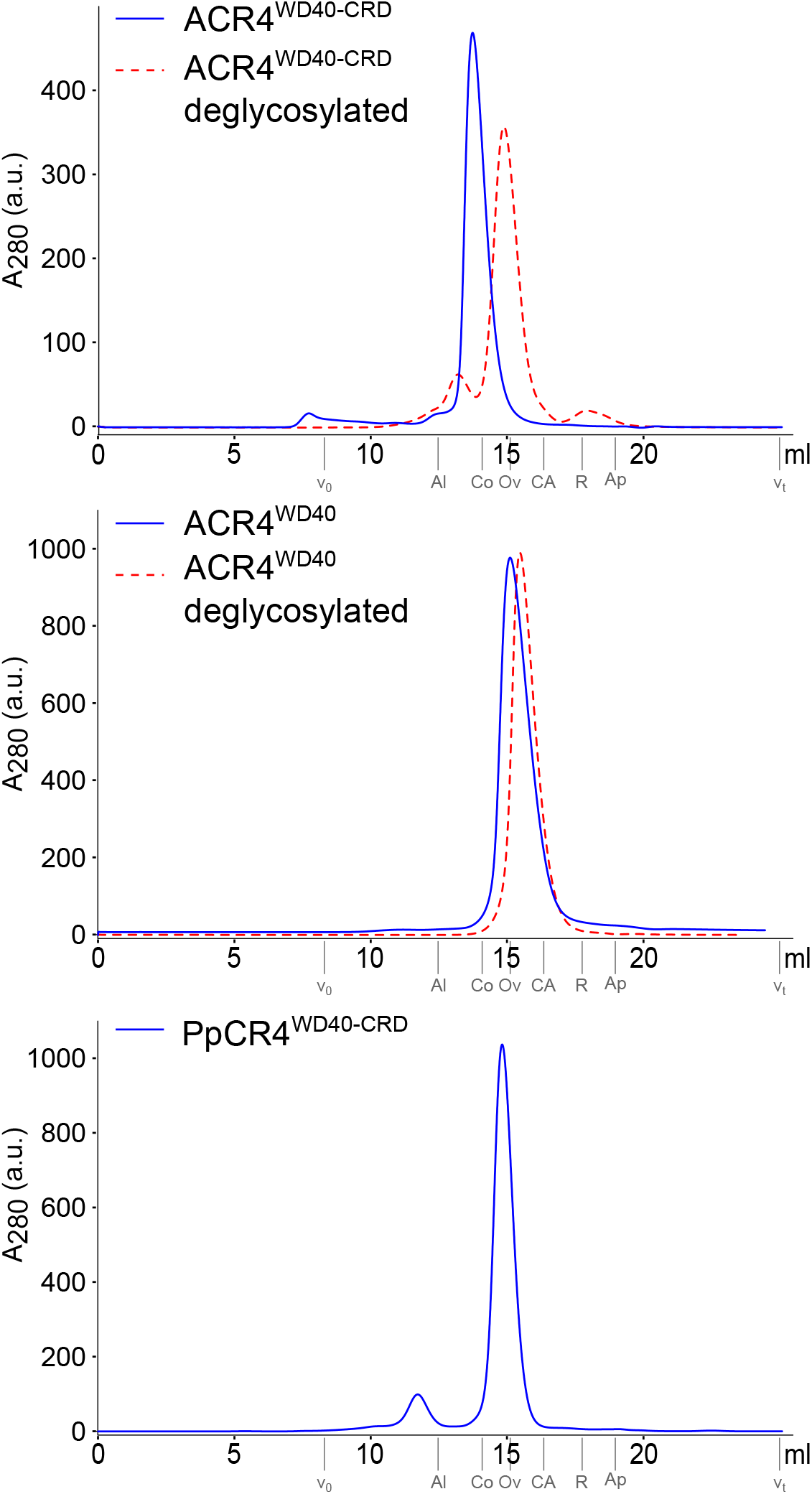
CRINKLY4 receptor ectodomains behave as monomers in solution. Analytical size-exclusion chromatography of the ACR4^WD40-CRD^, ACR4^WD40^ and PpCR4^WD40-CRD^ in the presence or absence of enzymatic deglycosylation. The void volume (V_0_), the total column volume (V_t_), and the elution volumes for molecular-mass standards (Al, Aldolase, 158 kDa; Co, Conalbumin, 75 kDa; Ov, Ovalbumin, 44 kDa; CA, Carbonic Anhydrase, 29 kDa; R, Ribonuclease A, 13.7 kDa; Ap, Aprotinin; 6.5 kDa) are indicated.

**Supplementary Figure 3.**
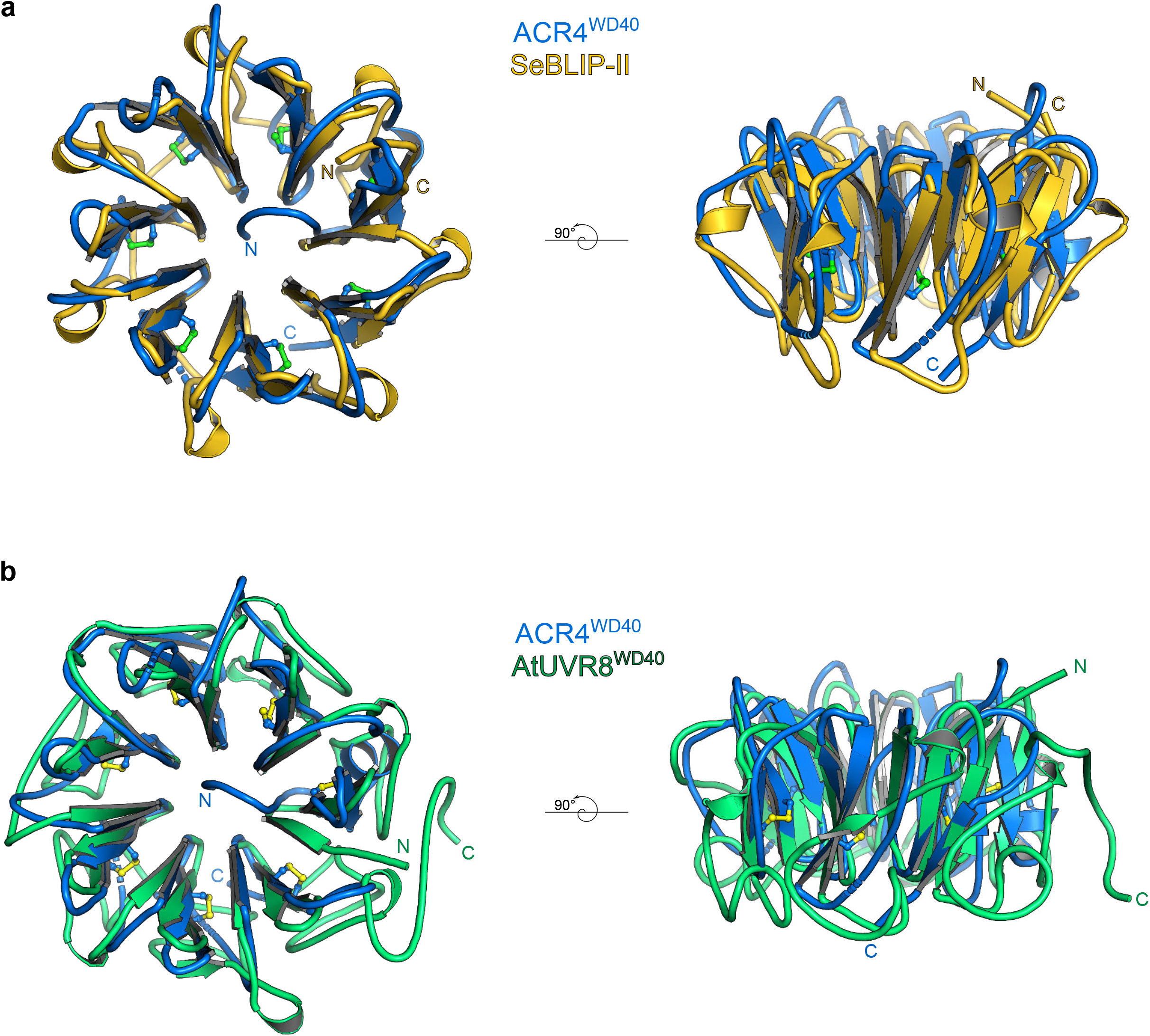
ACR4^WD40^ shares structural features with known WD40 domains. Structural superposition of ACR4^WD40^ (blue ribbon diagram) with **a**, the secreted β-lactamase inhibitor protein II BLIP-II (PDB-ID 1JTD^39^, in yellow) from the bacterium *Streptomyces exfoliatus* (r.m.s.d. is ∼2.2 Å comparing 192 corresponding C_α_ atoms), and **b**, with the WD40 domain of the UV-B photoreceptor UVR8 (PDB-ID 4D9S^40^, r.m.s.d. is ∼2.4 Å comparing 218 corresponding C_α_ atoms). Note that SeBLIP-II and UVR8 shares the blade number and overall architecture with ACR4^WD40^, but lack the buried N-terminal strand and the conserved disulfide bonds stabilizing each blade.

**Supplementary Figure 4.**
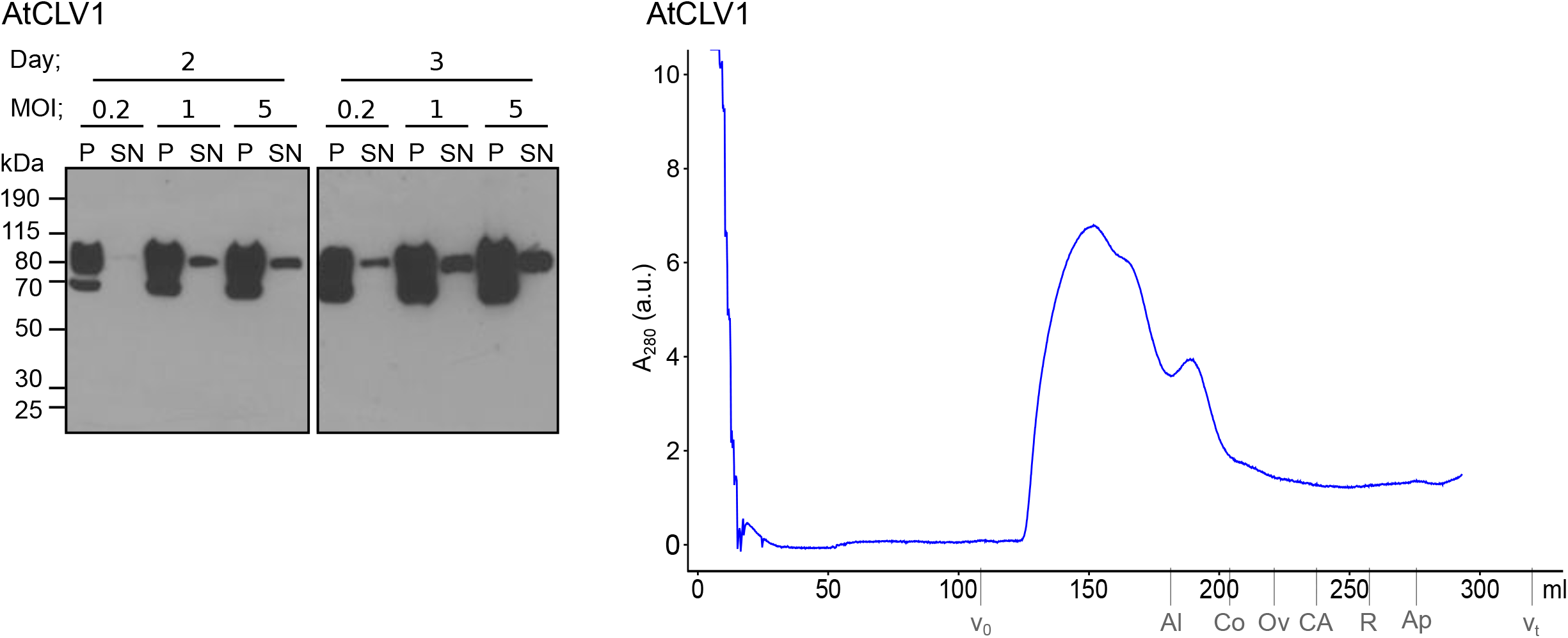
Expression and purification attempts of the AtCLV1 LRR ectodomain. Shown are immunoblot analyses monitoring the secreted expression of the AtCLV1 ectodomain (see Methods) with an anti-His antibody (left panels, Day, days post infection, MOI, multiplicity of infection; SN, supernatant; P, pellet). Right panel: Preparative size-exclusion chromatography of the purified AtCLV1 ectodomain reveals the presence of large aggregates. The void (V_0_), total (V_t_), and elution volumes for molecular-mass standards (Al, Aldolase, 158 kDa; Co, Conalbumin, 75 kDa; Ov, Ovalbumin, 44 kDa; CA, Carbonic Anhydrase, 29 kDa; R, Ribonuclease A, 13.7 kDa; Ap, Aprotinin; 6.5 kDa) are indicated.

**Supplementary Figure 5.**
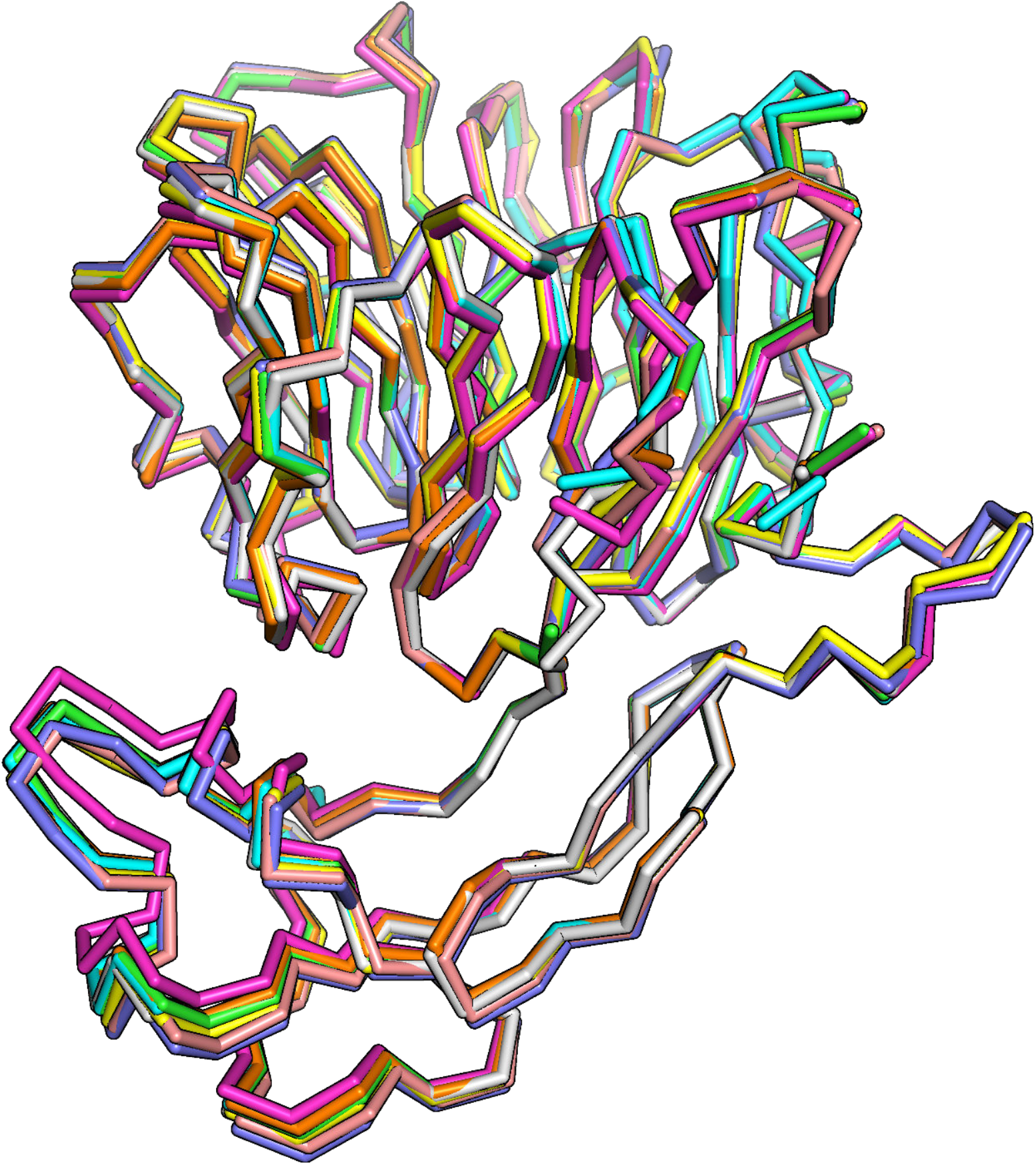
Only small WD40 - CRD inter-domain movements can be observed in the PpCR4 crystal structure. Structural superposition of the eight molecules located in the asymmetric unit of the PpCR4^WD40-CRD^ crystal structure (r.m.s.d. is ∼0.3-0.5 Å comparing 360 corresponding C_α_ atoms). Individual molecules are shown in different colors as C_α_ traces.

**Supplementary Figure 6.**
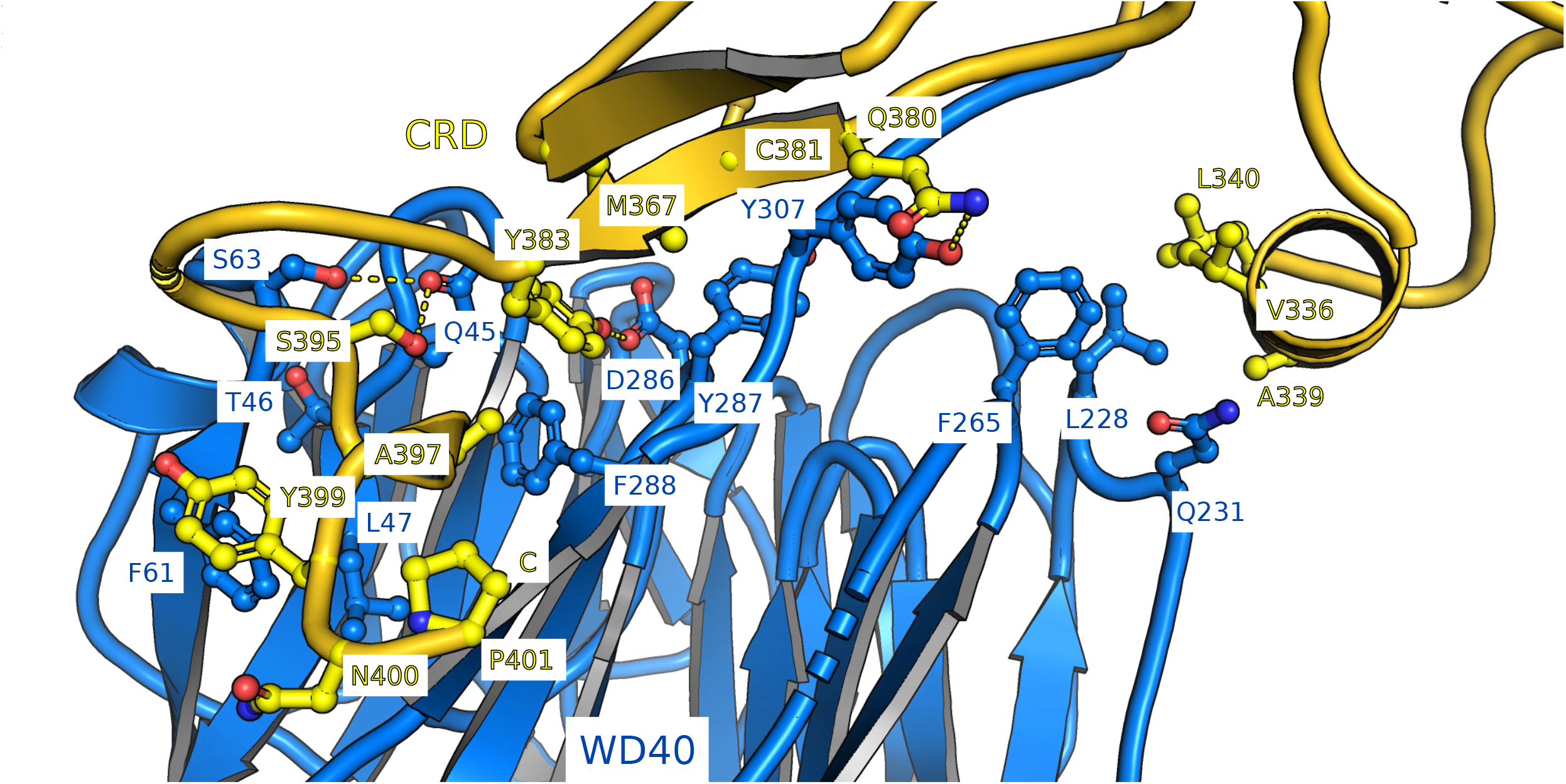
Overview of the WD40 – CRD domain interface in the PpCRD^WD40- CRD^ structure. Shown is a ribbon diagram of the PpCR4 ectodomain (colored according to Fig. 1e) with selected interface residues shown in bonds representation. Hydrogen bonds and salt bridges are indicated by dotted lines.

**Supplementary Figure 7.**
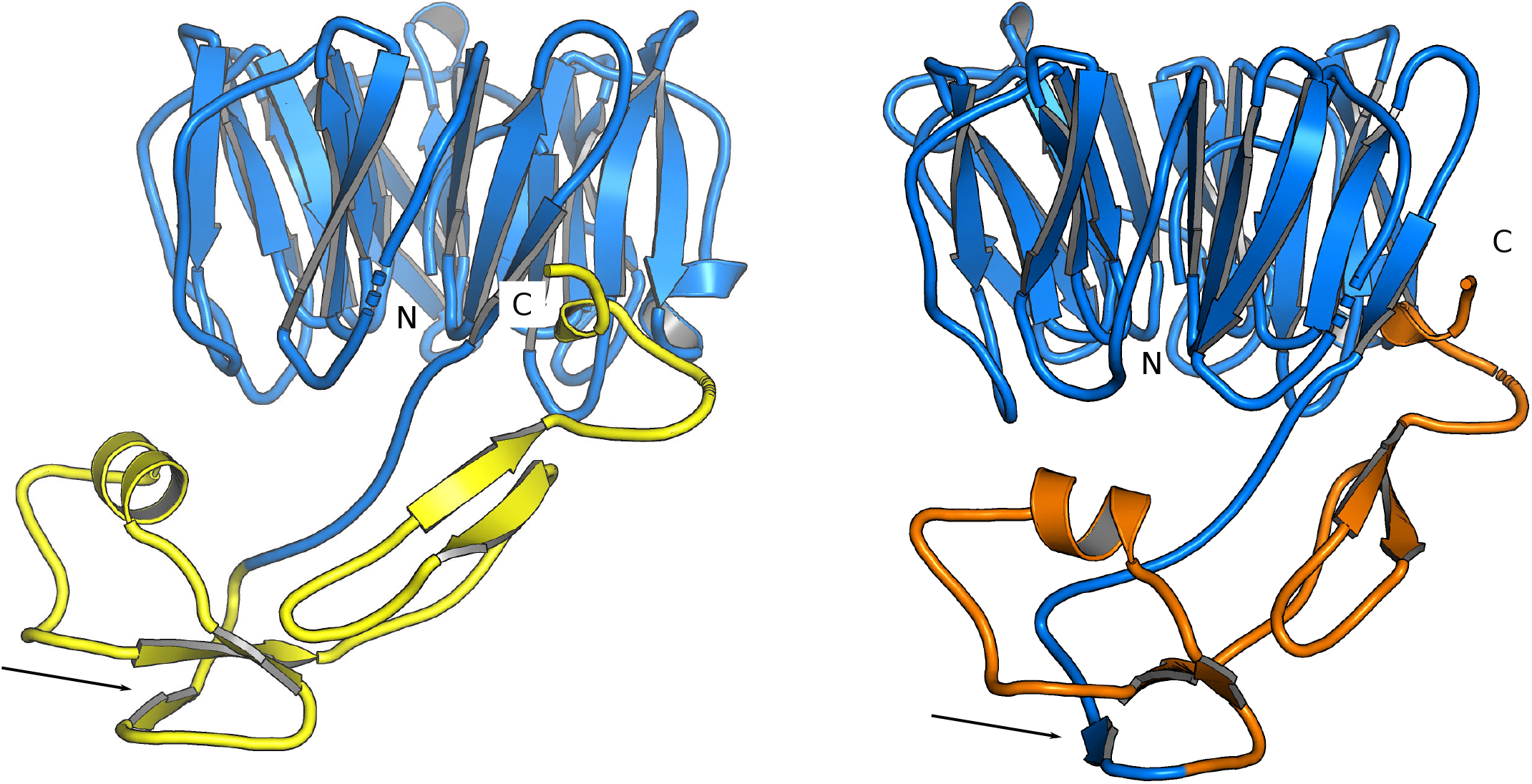
Structural visualization of the TNFR/CRD domain boundaries used in this and in previous studies. Ribbon diagram of PpCR4^WD40-CRD^ with the WD40 domain shown in blue and the experimentally determined CRD domain boundaries shown in yellow (left panel). The previously used TNFR domain boundaries^20,24^ derived from sequence analysis (in orange) omit the most N-terminal β-strand in the CRD (in blue, indicated by a black arrow).

**Supplementary Figure 8.**
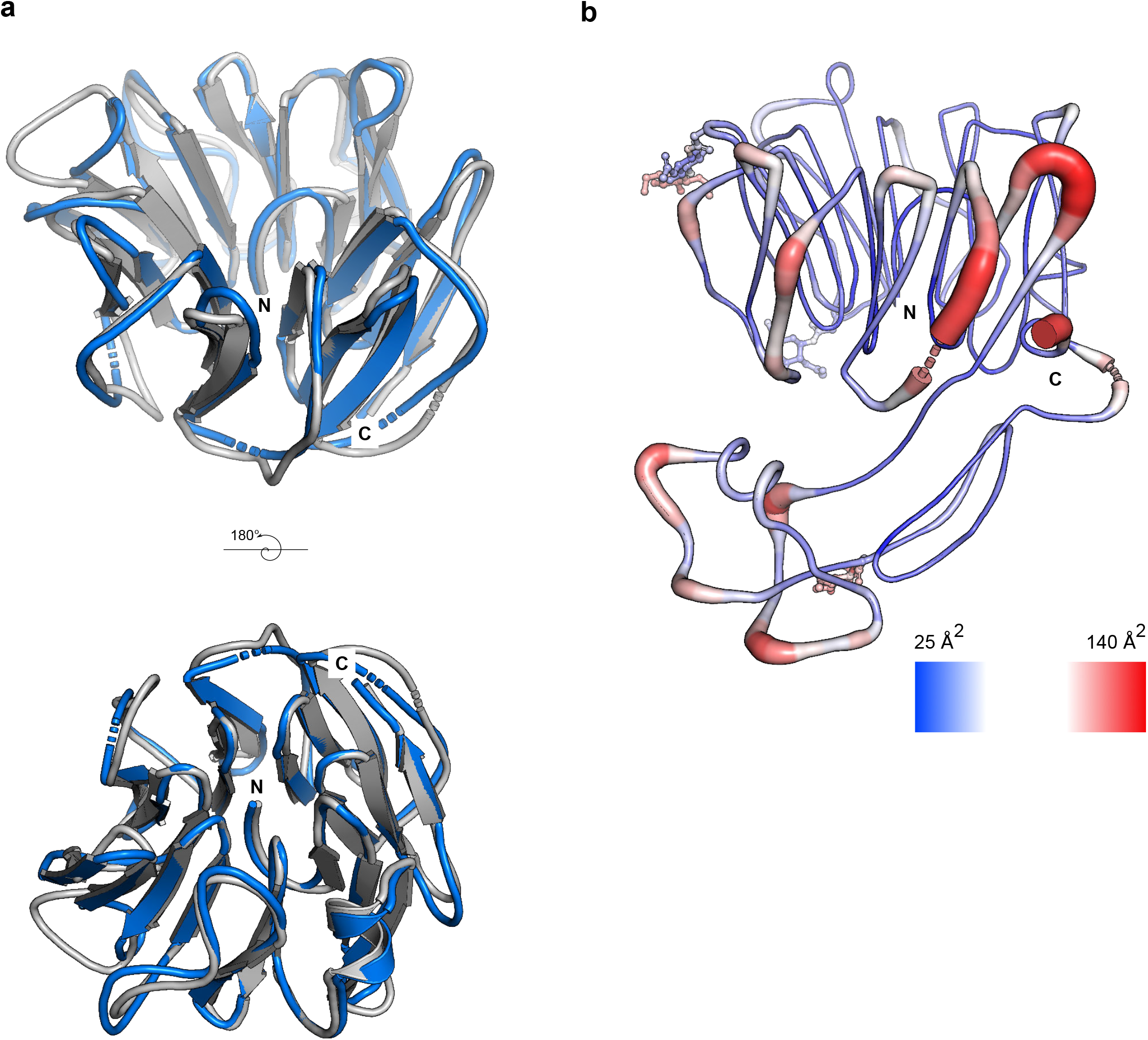
Structurally conserved loop regions contribute to the formation of a putative ligand binding groove in CRINKLY4 WD40 domains. **a**, Structural superposition of the isolated WD40 domain from ACR4 (blue) and PpCR4 (light gray, r.m.s.d. is is ∼1.4 Å comparing 246 corresponding C_α_ atoms reveals the loop regions contributing to the formation of a putative ligand binding groove to adopt similar orientations in both structures. **b**, A temperature (B-) factor plot of PpCR4^WD40-CRD^ (molecule chain A) reveals little structural flexibility for the secondary structure elements forming part of the putative binding groove, while the partially disordered loops connecting the blades of the β-propeller and the loops connecting the CRD appear mobile in the PpCR4^WD40-CRD^ crystal structure.

**Supplementary Table 1.**
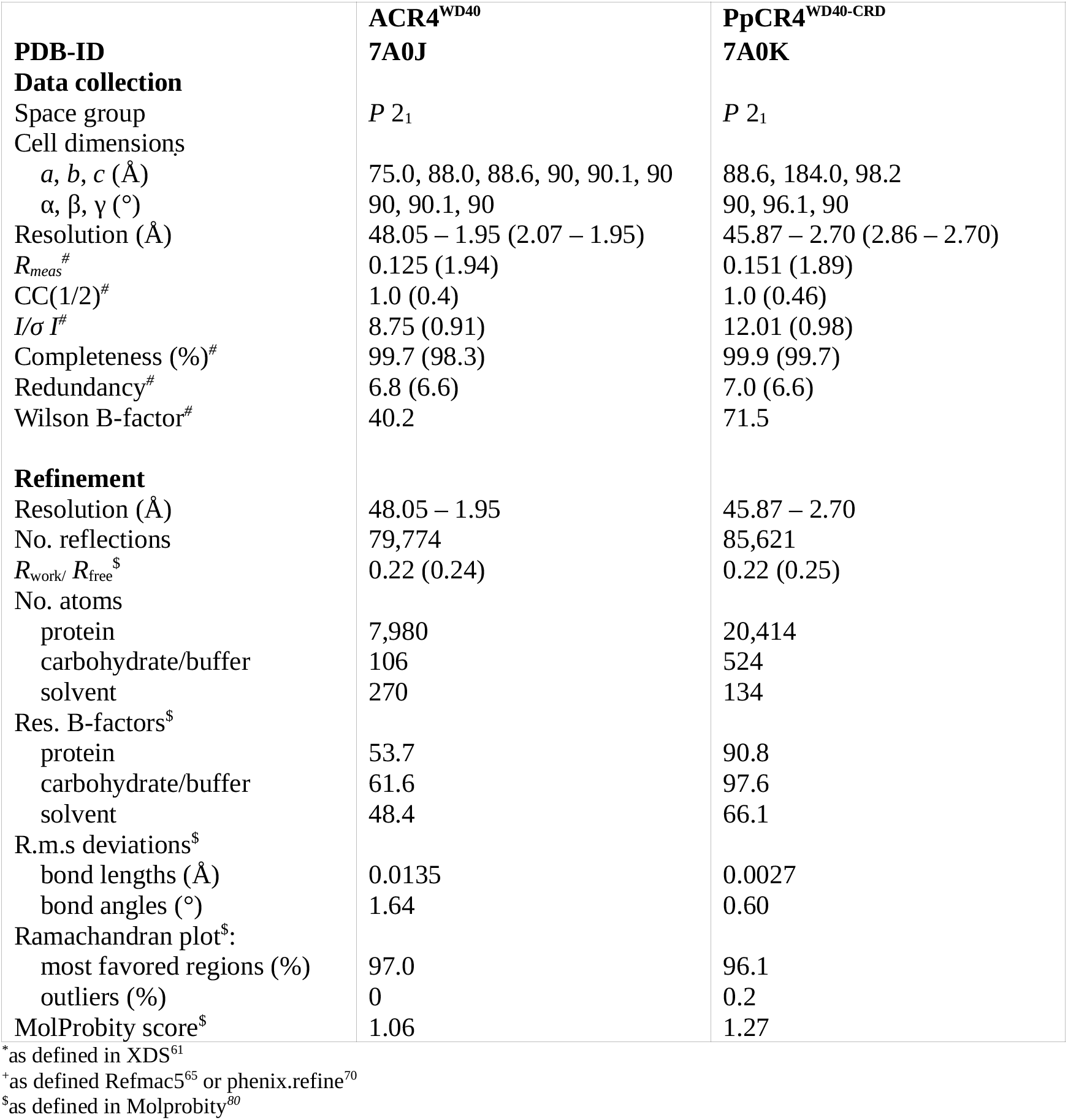
Crystallographic data collection and refinement statistics.

**Supplementary Table 2.**
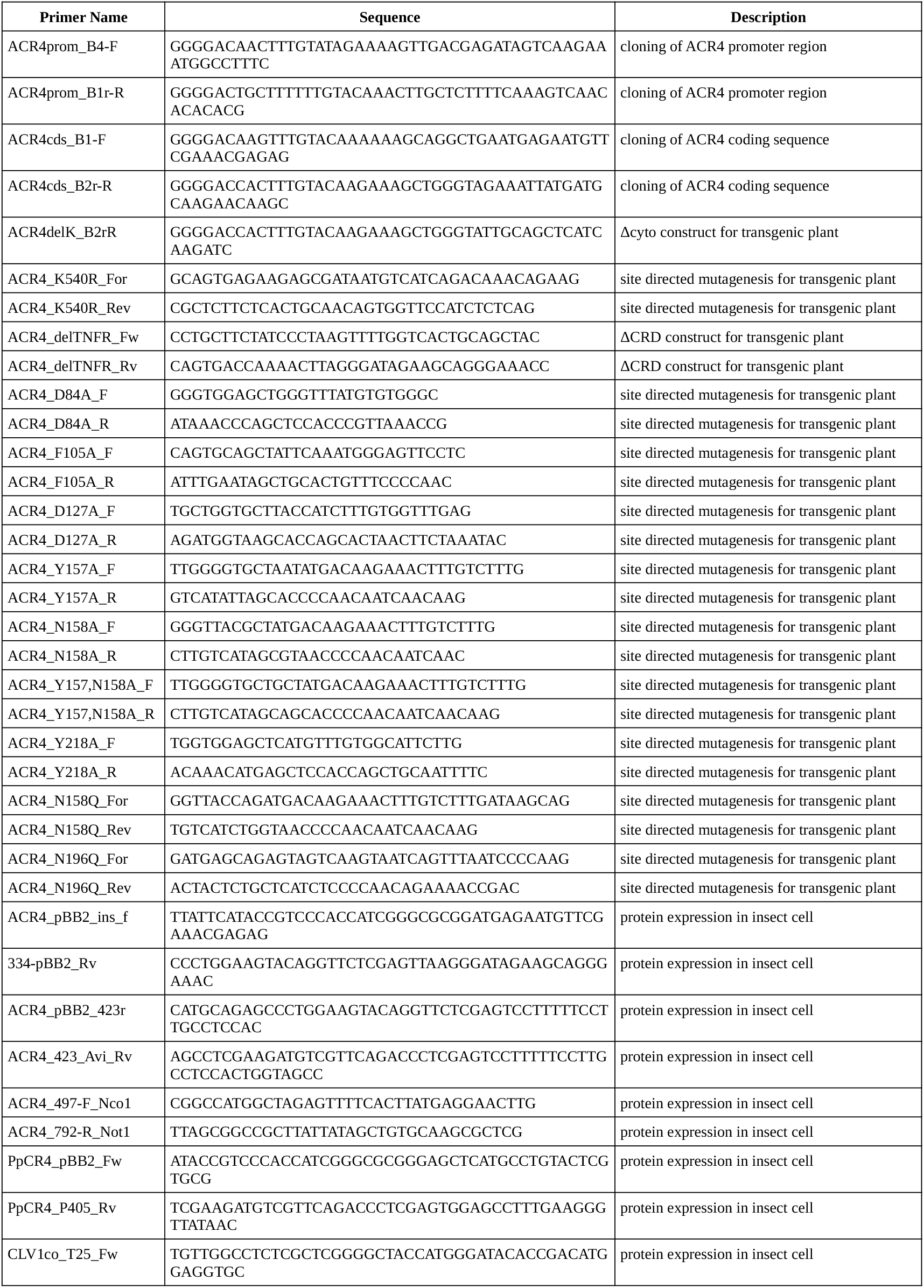
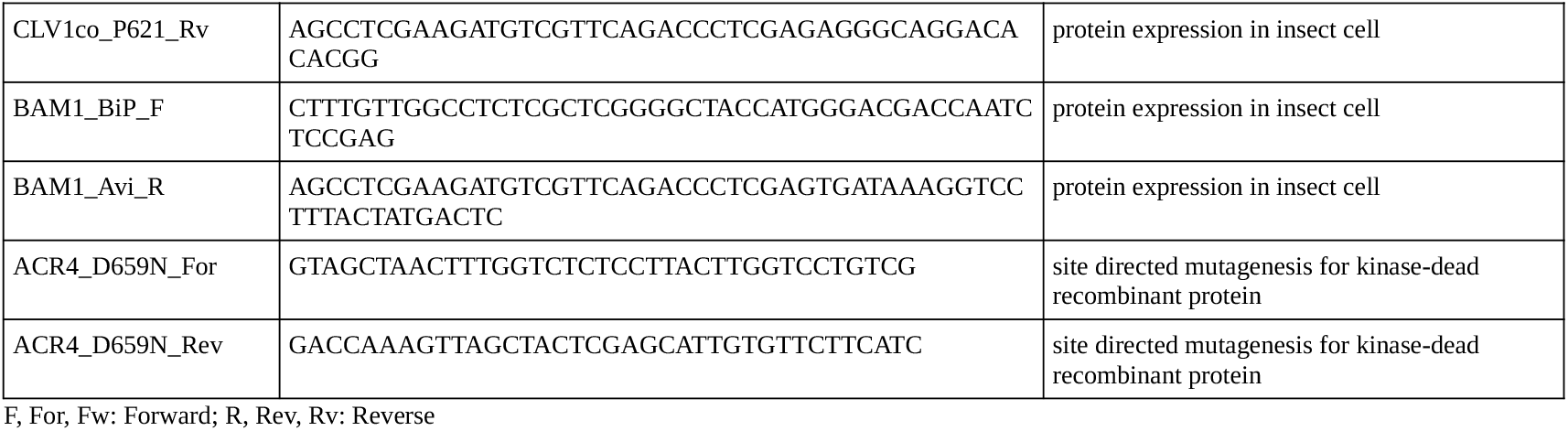
Primers used in this study

**Supplementary Table 3.**
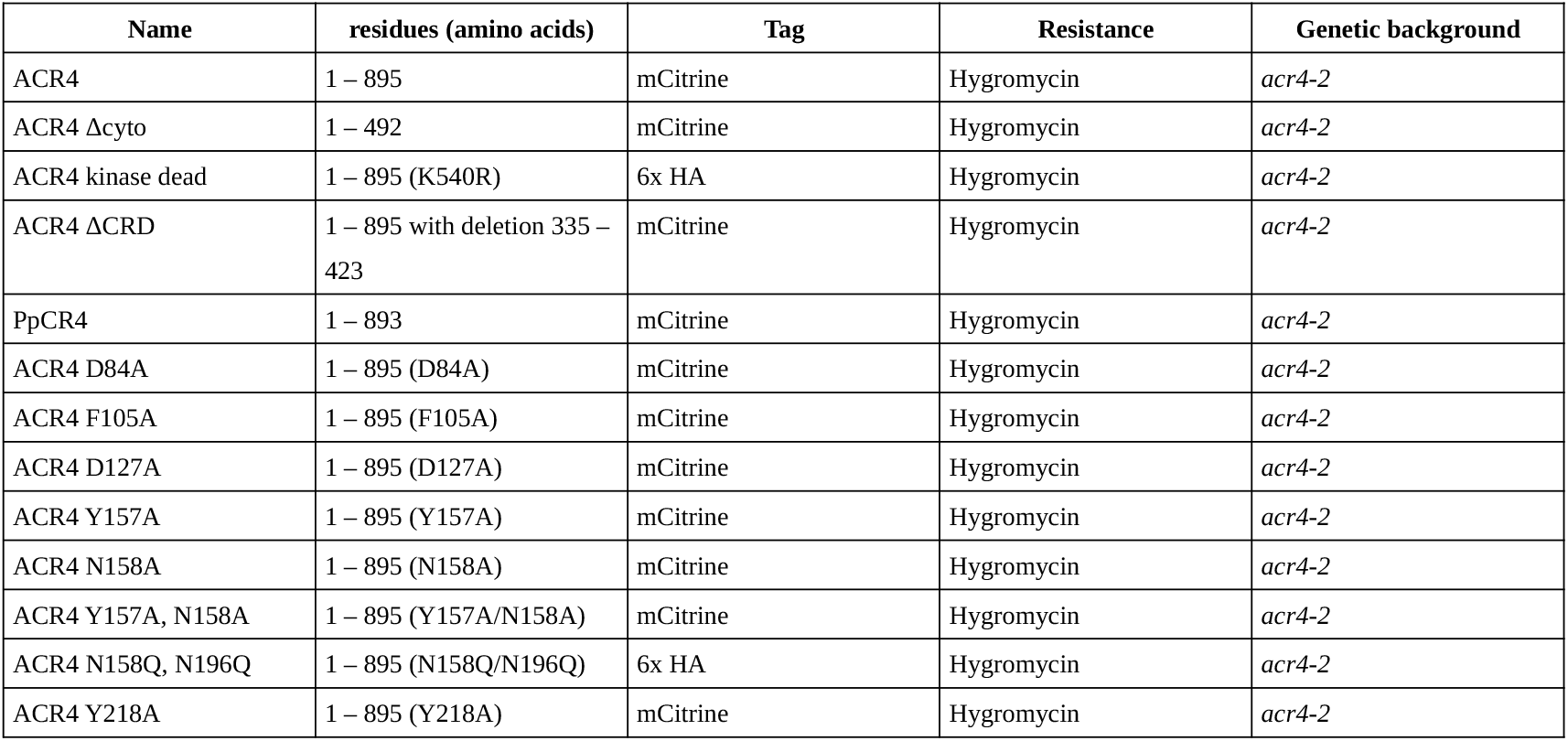
Transgenic lines generated in this study

| Name | residues (amino acids) | Tag | Resistance | Genetic background |
| --- | --- | --- | --- | --- |
| ACR4 | 1 – 895 | mCitrine | Hygromycin | *acr4-2* |
| ACR4 Δcyto | 1 – 492 | mCitrine | Hygromycin | *acr4-2* |
| ACR4 kinase dead | 1 – 895 (K540R) | 6x HA | Hygromycin | *acr4-2* |
| ACR4 ΔCRD | 1 – 895 with deletion 335 –<br>423 | mCitrine | Hygromycin | *acr4-2* |
| PpCR4 | 1 – 893 | mCitrine | Hygromycin | *acr4-2* |
| ACR4 D84A | 1 – 895 (D84A) | mCitrine | Hygromycin | *acr4-2* |
| ACR4 F105A | 1 – 895 (F105A) | mCitrine | Hygromycin | *acr4-2* |
| ACR4 D127A | 1 – 895 (D127A) | mCitrine | Hygromycin | *acr4-2* |
| ACR4 Y157A | 1 – 895 (Y157A) | mCitrine | Hygromycin | *acr4-2* |
| ACR4 N158A | 1 – 895 (N158A) | mCitrine | Hygromycin | *acr4-2* |
| ACR4 Y157A, N158A | 1 – 895 (Y157A/N158A) | mCitrine | Hygromycin | *acr4-2* |
| ACR4 N158Q, N196Q | 1 – 895 (N158Q/N196Q) | 6x HA | Hygromycin | *acr4-2* |
| ACR4 Y218A | 1 – 895 (Y218A) | mCitrine | Hygromycin | *acr4-2* |

## References

1. Shiu, S. H. & Bleecker, A. B. Receptor-like kinases from Arabidopsis form a monophyletic gene family related to animal receptor kinases. Proc. Natl. Acad. Sci. U. S. A. 98, 10763–10768 (2001).

2. Hohmann, U., Lau, K. & Hothorn, M. The Structural Basis of Ligand Perception and Signal Activation by Receptor Kinases. Annu. Rev. Plant Biol. 68, 109–137 (2017).

3. Bozsoki, Z. et al. Receptor-mediated chitin perception in legume roots is functionally separable from Nod factor perception. Proc. Natl. Acad. Sci. U. S. A. 114, E8118–E8127 (2017).

4. Liu, T. et al. Chitin-Induced Dimerization Activates a Plant Immune Receptor. Science 336, 1160–1164 (2012).

5. Liu, S. et al. Molecular Mechanism for Fungal Cell Wall Recognition by Rice Chitin Receptor OsCEBiP. Structure 24, 1192–1200 (2016).

6. Ma, R. et al. Structural basis for specific self-incompatibility response in Brassica. Cell Res. 26, 1320–1329 (2016).

7. Moussu, S. & Santiago, J. Structural biology of cell surface receptor-ligand interactions. Curr. Opin. Plant Biol. 52, 38–45 (2019).

8. Vaattovaara, A. et al. Mechanistic insights into the evolution of DUF26-containing proteins in land plants. Commun. Biol. 2, 56 (2019).

9. Moussu, S., Augustin, S., Roman, A. O., Broyart, C. & Santiago, J. Crystal structures of two tandem malectin-like receptor kinases involved in plant reproduction. Acta Crystallogr. Sect. Struct. Biol. 74, 671–680 (2018).

10. Xiao, Y. et al. Mechanisms of RALF peptide perception by a heterotypic receptor complex. Nature 572, 270–274 (2019).

11. Moussu, S. et al. Structural basis for recognition of RALF peptides by LRX proteins during pollen tube growth. Proc. Natl. Acad. Sci. U. S. A. 117, 7494–7503 (2020).

12. Becraft, P. W., Stinard, P. S. & McCarty, D. R. CRINKLY4: A TNFR-like receptor kinase involved in maize epidermal differentiation. Science 273, 1406–1409 (1996).

13. Jin, P., Guo, T. & Becraft, P. W. The maize CR4 receptor-like kinase mediates a growth factor-like differentiation response. Genesis 27, 104–116 (2000).

14. Banner, D. W. et al. Crystal structure of the soluble human 55 kd TNF receptor-human TNF beta complex: implications for TNF receptor activation. Cell 73, 431–445 (1993).

15. Sprang, S. R. The divergent receptors for TNF. Trends Biochem. Sci. 15, 366–368 (1990).

16. Becraft, P. W., Kang, S. H. & Suh, S. G. The maize CRINKLY4 receptor kinase controls a cell-autonomous differentiation response. Plant Physiol. 127, 486–496 (2001).

17. Tanaka, H. et al. ACR4, a putative receptor kinase gene of Arabidopsis thaliana, that is expressed in the outer cell layers of embryos and plants, is involved in proper embryogenesis. Plant Cell Physiol. 43, 419–428 (2002).

18. Gifford, M. L., Dean, S. & Ingram, G. C. The Arabidopsis ACR4 gene plays a role in cell layer organisation during ovule integument and sepal margin development. Development 130, 4249–4258 (2003).

19. Watanabe, M., Tanaka, H., Watanabe, D., Machida, C. & Machida, Y. The ACR4 receptor-like kinase is required for surface formation of epidermis-related tissues in Arabidopsis thaliana. Plant J. 39, 298–308 (2004).

20. Gifford, M. L., Robertson, F. C., Soares, D. C. & Ingram, G. C. ARABIDOPSIS CRINKLY4 function, internalization, and turnover are dependent on the extracellular crinkly repeat domain. Plant Cell 17, 1154–1166 (2005).

21. Cao, X., Li, K., Suh, S.-G., Guo, T. & Becraft, P. W. Molecular analysis of the CRINKLY4 gene family in Arabidopsis thaliana. Planta 220, 645–657 (2005).

22. Meyer, M. R., Lichti, C. F., Townsend, R. R. & Rao, A. G. Identification of in vitro autophosphorylation sites and effects of phosphorylation on the Arabidopsis CRINKLY4 (ACR4) receptor-like kinase intracellular domain: insights into conformation, oligomerization, and activity. Biochemistry 50, 2170–2186 (2011).

23. Zhao, Y., Liu, X., Xu, Z., Yang, H. & Li, J. Characterization and enzymatic properties of protein kinase ACR4 from Arabidopsis thaliana. Biochem. Biophys. Res. Commun. 489, 270–274 (2017).

24. Qin, Y. et al. Redox-Mediated Endocytosis of a Receptor-Like Kinase during Distal Stem Cell Differentiation Depends on Its Tumor Necrosis Factor Receptor Domain. Plant Physiol. 181, 1075–1095 (2019).

25. Nikonorova, N., Yue, K., Beeckman, T. & De Smet, I. Arabidopsis research requires a critical re-evaluation of genetic tools. J. Exp. Bot. 69, 3541–3544 (2018).

26. Roeder, A. H. K., Cunha, A., Ohno, C. K. & Meyerowitz, E. M. Cell cycle regulates cell type in the Arabidopsis sepal. Dev. Camb. Engl. 139, 4416–4427 (2012).

27. Xu, C. & Min, J. Structure and function of WD40 domain proteins. Protein Cell 2, 202–214 (2011).

28. De Smet, I. et al. Receptor-like kinase ACR4 restricts formative cell divisions in the Arabidopsis root. Science 322, 594–597 (2008).

29. Stahl, Y., Wink, R. H., Ingram, G. C. & Simon, R. A signaling module controlling the stem cell niche in Arabidopsis root meristems. Curr. Biol. 19, 909–914 (2009).

30. Stahl, Y. & Simon, R. Peptides and receptors controlling root development. Philos. Trans. R. Soc. Lond. B. Biol. Sci. 367, 1453–1460 (2012).

31. Berckmans, B., Kirschner, G., Gerlitz, N., Stadler, R. & Simon, R. CLE40 Signaling Regulates Root Stem Cell Fate. Plant Physiol. 182, 1776–1792 (2020).

32. Stahl, Y. et al. Moderation of Arabidopsis root stemness by CLAVATA1 and ARABIDOPSIS CRINKLY4 receptor kinase complexes. Curr. Biol. CB 23, 362–371 (2013).

33. Yue, K. et al. PP2A-3 interacts with ACR4 and regulates formative cell division in the Arabidopsis root. Proc. Natl. Acad. Sci. U. S. A. 113, 1447–1452 (2016).

34. Meyer, M. R., Shah, S., Zhang, J., Rohrs, H. & Rao, A. G. Evidence for intermolecular interactions between the intracellular domains of the arabidopsis receptor-like kinase ACR4, its homologs and the Wox5 transcription factor. PloS One 10, e0118861 (2015).

35. Demko, V., Ako, E., Perroud, P.-F., Quatrano, R. & Olsen, O.-A. The phenotype of the CRINKLY4 deletion mutant of Physcomitrella patens suggests a broad role in developmental regulation in early land plants. Planta 244, 275–284 (2016).

36. Nikonorova, N., Vu, L. D., Czyzewicz, N., Gevaert, K. & De Smet, I. A phylogenetic approach to study the origin and evolution of the CRINKLY4 family. Front. Plant Sci. 6, 880 (2015).

37. Krissinel, E. & Henrick, K. Inference of macromolecular assemblies from crystalline state. J. Mol. Biol. 372, 774–797 (2007).

38. Holm, L. & Sander, C. Protein structure comparison by alignment of distance matrices. J. Mol. Biol. 233, 123–138 (1993).

39. Lim, D. et al. Crystal structure and kinetic analysis of beta-lactamase inhibitor protein-II in complex with TEM-1 beta-lactamase. Nat. Struct. Biol. 8, 848–852 (2001).

40. Christie, J. M. et al. Plant UVR8 photoreceptor senses UV-B by tryptophan-mediated disruption of cross-dimer salt bridges. Science 335, 1492–1496 (2012).

41. Rizzini, L. et al. Perception of UV-B by the Arabidopsis UVR8 protein. Science 332, 103–106 (2011).

42. Pugalenthi, G., Nithya, V., Chou, K.-C. & Archunan, G. Nglyc: A Random Forest Method for Prediction of N-Glycosylation Sites in Eukaryotic Protein Sequence. Protein Pept. Lett. 27, 178–186 (2020).

43. Lau, K., Podolec, R., Chappuis, R., Ulm, R. & Hothorn, M. Plant photoreceptors and their signaling components compete for COP1 binding via VP peptide motifs. EMBO J. 38, e102140 (2019).

44. Anne, P. et al. CLERK is a novel receptor kinase required for sensing of root-active CLE peptides in Arabidopsis. Development 145, pii: dev162354 (2018).

45. Okuda, S. et al. Molecular mechanism for the recognition of sequence-divergent CIF peptides by the plant receptor kinases GSO1/SGN3 and GSO2. Proc. Natl. Acad. Sci. U. S. A. 117, 2693–2703 (2020).

46. Stokes, K. D. & Gururaj Rao, A. Dimerization properties of the transmembrane domains of Arabidopsis CRINKLY4 receptor-like kinase and homologs. Arch. Biochem. Biophys. 477, 219–226 (2008).

47. Stokes, K. D. & Rao, A. G. The role of individual amino acids in the dimerization of CR4 and ACR4 transmembrane domains. Arch. Biochem. Biophys. 502, 104–111 (2010).

48. Arabidopsis Genome Initiative. Analysis of the genome sequence of the flowering plant Arabidopsis thaliana. Nature 408, 796–815 (2000).

49. Rensing, S. A. et al. The Physcomitrella genome reveals evolutionary insights into the conquest of land by plants. Science 319, 64–69 (2008).

50. Haruta, M., Sabat, G., Stecker, K., Minkoff, B. B. & Sussman, M. R. A peptide hormone and its receptor protein kinase regulate plant cell expansion. Science 343, 408–411 (2014).

51. Kessler, S. A., Lindner, H., Jones, D. S. & Grossniklaus, U. Functional analysis of related CrRLK1L receptor-like kinases in pollen tube reception. EMBO Rep. 16, 107–115 (2015).

52. Haruta, M., Gaddameedi, V., Burch, H., Fernandez, D. & Sussman, M. R. Comparison of the effects of a kinase-dead mutation of FERONIA on ovule fertilization and root growth of Arabidopsis. FEBS Lett. 592, 2395–2402 (2018).

53. Chen, J. et al. FERONIA interacts with ABI2-type phosphatases to facilitate signaling cross-talk between abscisic acid and RALF peptide in Arabidopsis. Proc. Natl. Acad. Sci. U. S. A. 113, E5519–5527 (2016).

54. Franck, C. M. et al. The Protein Phosphatases ATUNIS1 and ATUNIS2 Regulate Cell Wall Integrity in Tip-Growing Cells. Plant Cell 30, 1906–1923 (2018).

55. Tanaka, H. et al. Novel receptor-like kinase ALE2 controls shoot development by specifying epidermis in Arabidopsis. Development 134, 1643–1652 (2007).

56. San-Bento, R., Farcot, E., Galletti, R., Creff, A. & Ingram, G. Epidermal identity is maintained by cell-cell communication via a universally active feedback loop in Arabidopsis thaliana. Plant J. 77, 46–58 (2014).

57. Cull, M. G. & Schatz, P. J. Biotinylation of proteins in vivo and in vitro using small peptide tags. Methods Enzymol. 326, 430–440 (2000).

58. Fairhead, M. & Howarth, M. Site-specific biotinylation of purified proteins using BirA. Methods Mol. Biol. 1266, 171–184 (2015).

59. Hashimoto, Y., Zhang, S. & Blissard, G. W. Ao38, a new cell line from eggs of the black witch moth, Ascalapha odorata (Lepidoptera: Noctuidae), is permissive for AcMNPV infection and produces high levels of recombinant proteins. BMC Biotechnol. 10, 50 (2010).

60. Bojar, D. et al. Crystal structures of the phosphorylated BRI1 kinase domain and implications for brassinosteroid signal initiation. Plant J. 78, 31–43 (2014).

61. Kabsch, W. Automatic processing of rotation diffraction data from crystals of initially unknown symmetry and cell constants. J. Appl. Crystallogr. 26, 795–800 (1993).

62. Sheldrick, G. M. A short history of SHELX. Acta Crystallogr. A 64, 112–122 (2008).

63. Bricogne, G., Vonrhein, C., Flensburg, C., Schiltz, M. & Paciorek, W. Generation, representation and flow of phase information in structure determination: recent developments in and around SHARP 2.0. Acta Crystallogr. D Biol. Crystallogr. 59, 2023–2030 (2003).

64. Terwilliger, T. C. et al. Iterative model building, structure refinement and density modification with the PHENIX AutoBuild wizard. Acta Crystallogr. D Biol. Crystallogr. 64, 61–69 (2008).

65. Murshudov, G. N., Vagin, A. A. & Dodson, E. J. Refinement of macromolecular structures by the maximum-likelihood method. Acta Crystallogr. D Biol. Crystallogr. 53, 240–255 (1997).

66. Evans, P. Scaling and assessment of data quality. Acta Crystallogr. D Biol. Crystallogr. 62, 72–82 (2006).

67. Lebedev, A. A. & Isupov, M. N. Space-group and origin ambiguity in macromolecular structures with pseudo-symmetry and its treatment with the program Zanuda. Acta Crystallogr. D Biol. Crystallogr. 70, 2430–2443 (2014).

68. McCoy, A. J. et al. Phaser crystallographic software. J. Appl. Crystallogr. 40, 658–674 (2007).

69. Emsley, P. & Cowtan, K. Coot: model-building tools for molecular graphics. Acta Crystallogr. D Biol. Crystallogr. 60, 2126–2132 (2004).

70. Afonine, P. V. et al. Towards automated crystallographic structure refinement with phenix.refine. Acta Crystallogr. D Biol. Crystallogr. 68, 352–367 (2012).

71. Goddard, T. D. et al. UCSF ChimeraX: Meeting modern challenges in visualization and analysis. Protein Sci. 27, 14–25 (2018).

72. Karimi, M., De Meyer, B. & Hilson, P. Modular cloning in plant cells. Trends Plant Sci. 10, 103–105 (2005).

73. Clough, S. J. & Bent, A. F. Floral dip: a simplified method forAgrobacterium-mediated transformation ofArabidopsis thaliana. Plant J. 16, 735–743 (1998).

74. Dunnett, C. W. A Multiple Comparison Procedure for Comparing Several Treatments with a Control. J. Am. Stat. Assoc. 50, 1096–1121 (1955).

75. Siegfried, S. & Hothorn, T. Count transformation models. Methods Ecol. Evol. 11, 818–827 (2020).

76. Hothorn, T., Bretz, F. & Westfall, P. Simultaneous inference in general parametric models. Biom. J. 50, 346–363 (2008).

77. Naismith, J. H., Devine, T. Q., Brandhuber, B. J. & Sprang, S. R. Crystallographic evidence for dimerization of unliganded tumor necrosis factor receptor. J. Biol. Chem. 270, 13303–13307 (1995).

78. Notredame, C., Higgins, D. G. & Heringa, J. T-coffee: a novel method for fast and accurate multiple sequence alignment1. J. Mol. Biol. 302, 205–217 (2000).

79. Kabsch, W. & Sander, C. Dictionary of protein secondary structure: pattern recognition of hydrogen-bonded and geometrical features. Biopolymers 22, 2577–2637 (1983).

80. Davis, I. W. et al. MolProbity: all-atom contacts and structure validation for proteins and nucleic acids. Nucleic Acids Res. 35, W375–383 (2007).

